# A yeast two-hybrid system to obtain triple-helical ligands from combinatorial random peptide libraries

**DOI:** 10.1101/2023.11.14.567114

**Authors:** Ryo Masuda, Khine Phyu Phyu Thant, Kazuki Kawahara, Hiroya Oki, Tetsuya Kadonosono, Yuji Kobayashi, Takaki Koide

**Author notes:** Corresponding author: Professor Takaki Koide, Ph.D., Department of Chemistry and Biochemistry, School of Advanced Science and Engineering, Waseda University, Shinjuku, Tokyo 169-8555, Japan. These authors contributed equally to this work (Ryo Masuda, Khine Phyu Phyu Thant).

## Abstract

Many bioactive proteins interact with collagen, recognizing amino acid sequences displayed on the triple helix. We report here a selection strategy to obtain triple-helical peptides that interact with the proteins from a combinatorial random library constructed in yeast cells. This system enables us to select them using the standard two-hybrid protocol, detecting interactions between triple-helical peptides and target proteins fused to the GAL4-activating and binding domains, respectively. The library was constructed having triple-helical peptides with a “host-guest” design in which host helix-stabilizing regions flanked guest random sequences. Using this system, we selected peptides that bind to pigment epithelium-derived factor (PEDF), a collagen-binding protein that shows anti-angiogenic and neurotrophic activities, from the libraries. Two-step selections from the total random library and subsequently from the second focused library yielded new PEDF-binding sequences that exhibited a comparable affinity to or more potent than that of the native PEDF-binding sequence in collagen. The obtained sequences also contained a variant of the PEDF-binding motif that did not match the known motif identified from the native collagen sequences. This combinatorial library system allows the chemical space of triple-helical peptides to be screened more widely than that found in native collagen, thus increasing the expectation of obtaining more specific and high-affinity peptides.

## Introduction

The triple helix, an iconic tertiary structure of proteins in the collagen family, is a right-handed supercoil of three left-handed polyproline-II helices consisting of tandem Gly-Xaa-Yaa tripeptide repeats. Pro residues at Xaa positions and post-translationally modified 4-hydroxyproline (Hyp) residues at Yaa positions are frequently found in natural collagen sequences, contributing to conformational thermal stability (1–3).

Fibril-forming types of collagen, such as types I, II, III, V, and XI, have long (>1000 amino acids) uninterrupted triple helices flanked by short telopeptides at both ends of the helices. These collagen types are prominent components of the extracellular matrix and play pivotal roles in maintaining the integrity of the organs and tissues of animals by forming rigid fibers. They also exhibit various physiological functions to regulate cell attachment, differentiation, angiogenesis, and blood coagulation (4,5). Such functions are elicited through specific interactions of bioactive macromolecules with certain regions in the triple helices. In many cases, the interactions are amino acid sequence-specific: collagen-binding integrins such as α1β1, α2β1, α10β1, and α11β1 recognize Gly-Phe-Hyp-Gly-Glu-Arg, Gly-Leu-Hyp-Gly-Glu-Asn, and other similar sequences containing Glu residues at Xaa positions (6–9). Discoidin-domain receptors (DDRs) 1 and 2, collagen-specific tyrosine-kinase receptors, recognize Gly-Val-Met-Gly-Phe-Hyp-containing sequences. The same region is shared by other collagen-binding proteins such as von Willebrand factor (VWF), a serum blood coagulation factor, and secreted protein acidic and rich in cysteine (SPARC), also known as osteonectin and BM-40, a factor regulating collagen assembly (10–13). Pigment epithelium-derived factor (PEDF), a neurotrophic/anti-angiogenic serpin, binds to the Lys-Gly-His-Arg-Gly-Phe-Ser-Gly-Leu sequence [named α1(I)930–938] in the type I collagen triple helix, which overlaps with the heparan sulfate proteoglycan (HSPG)-binding sites and crosslinking sites with the allysine-containing telopeptide region between different triple helices (14–17). Because two or more proteins share overlapping binding sites on the native collagen triple helix, the collagen triple helix performs complex regulation of physiological events with multiple intertwined macromolecular interactions.

Synthetic triple-helical peptides containing certain parts of natural collagen sequences have been effectively used to identify amino acid sequences in the collagen triple helices recognized by the collagen-binding macromolecules. Because most fragments of natural collagen generated by enzymatic or chemical cleavage lose the ability to maintain the triple-helical conformation, synthetic peptides are designed to be so-called “host-guest peptides” in which the natural collagen sequences are flanked by several repeats of Pro-Pro-Gly or Pro-Hyp-Gly that enhance the thermal stability of the triple helix (18,19). Farndale and co-workers established the collections of host-guest peptides that cover the entire amino acid sequences of fibril-forming types of collagen, and named them “Collagen Toolkits.” The amino acid sequences responsible for the binding to the collagen-binding integrins, VWF, DDRs, and SPARC (described above), were identified by the screening of Collagen Toolkits II and III, which covered the amino acid sequences of α1(II) and α1(III) chains, respectively (10–13). The PEDF-binding sequence, α1(I)930–938, was also identified by Koide and co-workers using synthetic triple-helical peptides (15).

Generally, the searching of random peptide libraries is a strategy to identify binding sequences for proteins of interest (POIs). Combinatorial random peptide libraries are constructed chemically by split-and-mix solid-phase synthesis or biologically by a DNA-encoded method involving degenerated codons (20,21). In principle, random peptide libraries contain a greater variety of peptides, including novel peptides that are not encoded in genomes. Such libraries, combined with high-throughput screening methods, are effective at obtaining a set of amino acid sequences recognized by POIs and determining the responsible binding sequence motifs.

In 2000, our group reported a yeast two-hybrid (Y2H) system to detect the interaction of peptides with heat-shock protein 47 (HSP47), a collagen-specific molecular chaperone (22–24). In that work, we constructed two collagen-like peptide libraries in which either six Xaa or Yaa positions of (Gly-Pro-Pro)_6_ were randomized in the form of the GAL4-activation domain (GAL4-AD) fusion proteins. The libraries were selected by the interaction with HSP47 fused to the GAL4-binding domain (GAL4-BD). Although the Xaa-randomized library yielded no hit clones, the selection enriched triple helix-stabilizing amino acid residues in the Yaa positions, showing a binding preference of HSP47 to the triple-helical conformation.

Here, we upgraded the design of libraries, in which both Xaa and Yaa positions of the Gly-Xaa-Yaa triplets were randomized to obtain triple-helical amino acid motifs recognized by POIs. The length of the randomized region was set based on the sizes of the binding interfaces of collagen-binding proteins reported to date. We also employed the host-guest design to increase the population of triple-helical peptides in the libraries. In the following sections, we describe the design, construction, and validation of the libraries, followed by the results of actual selections of PEDF-binding triple-helical peptides.

## Results

### Design, construction, and validation of the random peptide library for Y2H selection

To detect the interaction of a POI and a triple-helical peptide, we designed a yeast two-hybrid system, as shown in Fig. 1A. Here, a POI and a peptide with collagenous Gly-Xaa-Yaa (here, we used Yaa-Gly-Xaa repeats for experimental convenience) are expressed as fusion proteins of GAL4-BD and GAL4-AD, respectively. The AD-fusion proteins were expected to trimerize and concomitant with triple-helix formation. The trimer binds to one, two, or three molecules of the BD-fusion POI. The resulting oligomeric complex can activate the expression of the reporter genes as well as the general 1:1 complex (22).

**Fig. 1.**
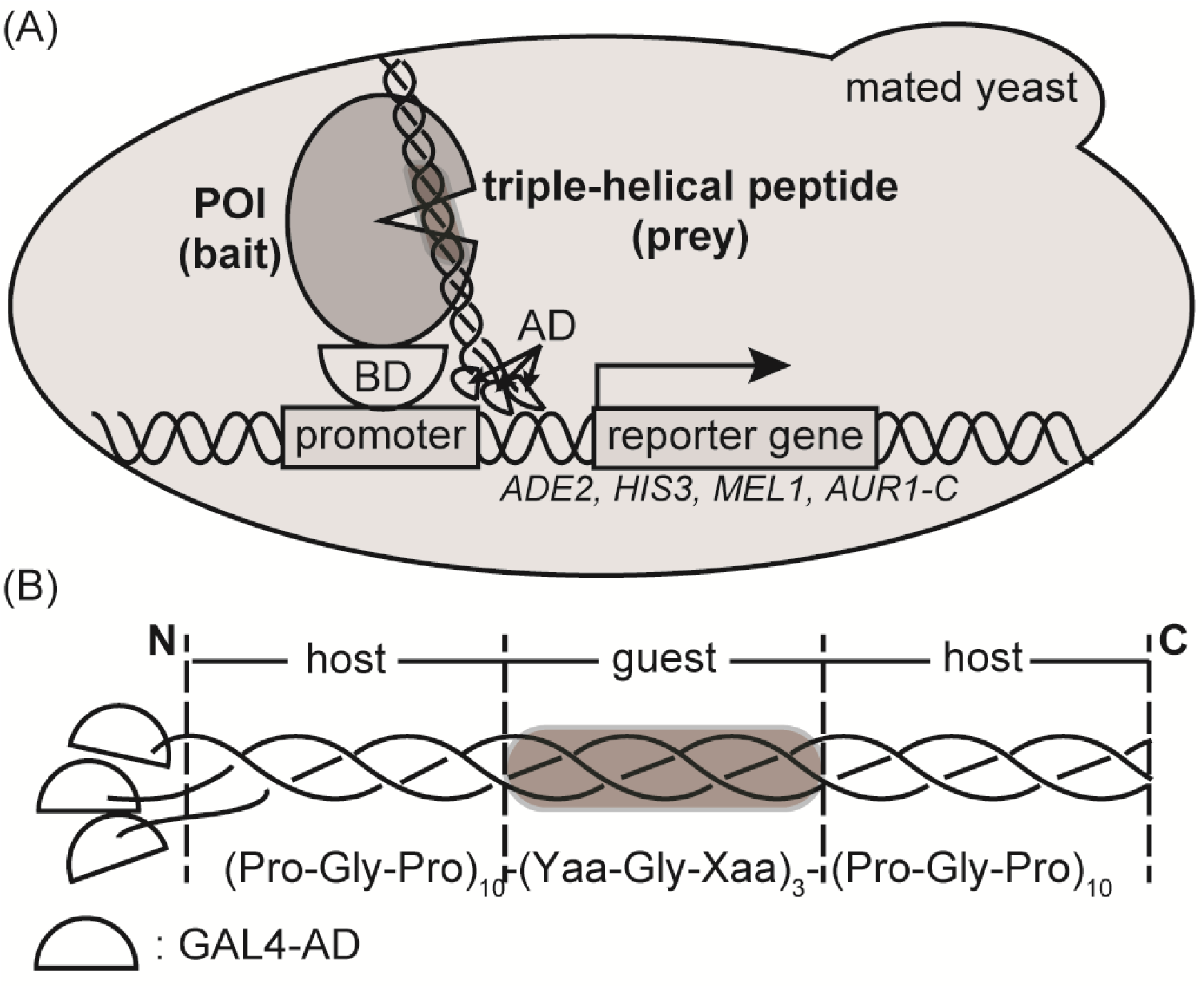
Design of the two-hybrid system for selecting protein-binding triple-helical peptides in yeast cells. (A) Mechanism of the two-hybrid assay detecting the binding between a protein of interest (POI) and a triple-helical peptide in a yeast cell. The interaction brings GAL4-AD and GAL4-BD into proximity, leading to expression of the reporter genes. (B) A construct of a GAL4-AD-fusion triple-helical peptide with a random region. Xaa and Yaa are randomized by setting the corresponding codons to NNK codons.

Fig. 1B shows the design of GAL4-AD-fusion random peptides with a propensity to form a triple helix. In the construct, we took advantage of the host-guest peptide for collagen-like triple-helical peptides (18,19). A middle portion consisting of three Yaa-Gly-Xaa repeats (guest) was flanked at each end by ten repeats of Pro-Gly-Pro triplet (host) with the ability to stabilize triple helices. NNK codons encoded each of three Xaa and Yaa amino acids to generate a randomized guest sequence in the peptides (N and K are equimolar mixtures of A/T/G/C and T/G, respectively). We randomized the three triplet repeats as the guest region, Yaa1-Gly2-Xaa3-Yaa4-Gly5-Xaa6-Yaa7-Gly8-Xaa9, because most collagen-binding proteins or domains recognize fewer than three triplets (9 amino acids), except the longer one (10 amino acids) for the catalytic domain of matrix metalloprotease-1 (Table S4, 6-15, 25). In theory, the diversity of the library is 20^6^ (= 6.4 × 10^7^). After the transformation of library plasmids into yeast cells, the diversity of the constructed library was estimated to be 4.0 × 10^6^, by counting the colony numbers.

The actual diversity of the random peptide library expressed in yeast cells was estimated by next-generation sequencing (NGS). The relative frequency of occurrences of Xaa and Yaa amino acid residues is shown in Fig. 2A. All possible Xaa-Yaa combinations were found in the library, and the frequencies of occurrence were similar to the theoretically expected ones (Fig. S1). The frequencies of Xaa-Yaa combinations in α-chains of human fibrillar collagen, α1(I), α2(I), α1(II), α1(III), α1(V), α2(V), α3(V), α1(XI), and α2(XI), are similarly shown for comparison (Fig. 2B). In contrast to the results of the peptide library, the frequency is significantly biased to the triple-helix-stabilizing Xaa-Yaa combinations. The Pro-Hyp combination with the highest triple-helix-stabilizing effect is the most prominent (11.2%) in human fibrillar collagen. Meanwhile, the frequency of occurrence decreases as the helix-stabilizing effect decreases. Notably, the combinations categorized with the lowest stabilization score do not occur (the combinations shown in blue in Fig. 2B). These findings indicate that the random peptide library, at least at the genetic level, contains a much greater variety of Xaa-Yaa combinations than human fibrillar collagen, including ones that had been rejected by natural selection.

**Fig. 2.**
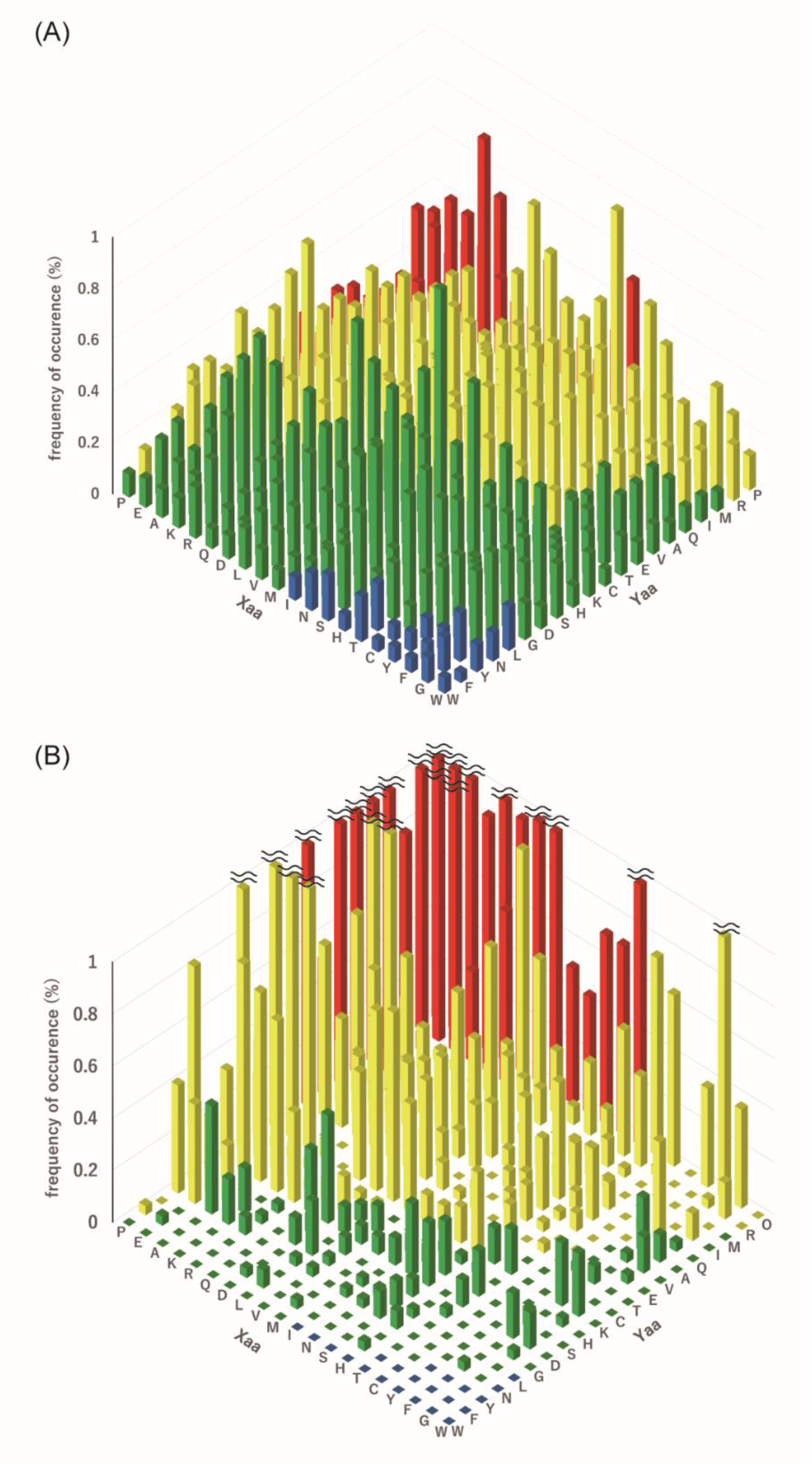
Distribution of Gly-Xaa-Yaa triplets in (A) the guest region of the yeast library and (B) the triple-helical region of human fibrillar collagen. Each triplet is categorized into four groups according to its predicted or experimentally observed effect on the thermostability of the triple helix, as previously reported (19). Here, the effect on the thermal stability, Δ*T* ^GPO^, is defined as the difference of melting temperature of the triple helix (*T*_m_) between (Gly-Pro-Hyp)_8_ and peptides in which a Gly-Pro-Hyp triplet is replaced with a corresponding Gly-Xaa-Yaa: Red: −10°C<Δ*T*_m_^GPO^, yellow: −20°C<Δ*T*_m_^GPO^<−10°C, green: −30°C<Δ*T*_m_^GPO^<−20°C, and blue Δ*T*_m_^GPO^<−30°C. “O” denotes Hyp. A reduced view of Fig. 2B is shown in Fig. S1B.

### Triple-helix-forming propensity of the peptides in yeast cells

The molecular chaperone HSP47 selectively binds to collagen-like peptides with a triple-helical conformation (22,26). Although the high-affinity binding site contains an Arg at the Yaa position (27), the triple-helical Pro-Pro-Gly repeating sequence is also recognized by HSP47 (28). This structural requirement of HSP47 for recognizing the repeating sequence is useful for determining the conformational states of the peptides expressed in yeast cells. To validate the lengths of the collagen-like peptides in the library and set the temperature at which Y2H selection could occur, the binding of HSP47 to (Pro-Gly-Pro)-repeating peptides with several repeating numbers was first examined.

We expressed (Pro-Gly-Pro)-repeating peptides with a repeat number of 10, 13, or 23, as GAL4-AD-fusion proteins in yeast cells. The lengths of (Pro-Gly-Pro)_10_, (Pro-Gly-Pro)_13_, and (Pro-Gly-Pro)_23_ correspond to those of the *N*-terminal host, the guest plus the *N*-terminal host, and the guest plus the *N*-and *C*-hosts in the above peptide design (Fig. 1B), respectively. The peptides with fewer (Pro-Gly-Pro)-repeats have lower thermal stability of the triple helices (19). We set three different culture temperatures, 16, 30, and 37°C, with the expectation of identifying the binding of BD-fusion HSP47 to these three peptides with GAL4-AD by the yeast two-hybrid system (Fig. 3). The mated yeast cells were selected by His and Ade auxotrophy in the presence of aureobasidin and X-α-Gal. When HSP47 interacts with the peptides, GAL4-BD and GAL4-AD are brought into proximity to activate the transcription of reporter genes (*AUR1-C*, *ADE2*, *HIS3*, and *MEL1*). The reporter gene expression allows yeast cells to survive on the plates and turn blue by degrading X-α-Gal. The interaction between HSP47 (bait) and peptides (prey) was detected by the growth of blue colonies on the selection plates. Overall, culturing at lower temperatures resulted in the growth of blue colonies expressing the peptides with shorter Pro-Gly-Pro repeats. HSP47 bound only to (Pro-Gly-Pro)_23_ in yeast cells at 37°C. At 30°C, HSP47 bound to (Pro-Gly-Pro)_13_ as well as (Pro-Gly-Pro)_23_, but not to (Pro-Gly-Pro)_10_. At 16°C, HSP47 bound to all three of the peptides. This temperature-dependent result demonstrated that HSP47 distinguished the conformational state of the host Pro-Gly-Pro repeats in yeast cells. Since the marked decrease in the colony growth expressing (Pro-Gly-Pro)_23_ was observed at 30°C, the library screening should be conducted below the temperature. It also indicates that BD-fusion HSP47 can be used as an indicator for determining the triple-helical conformation of peptides expressed in yeast cells because it binds to (Pro-Gly-Pro) repeats in the host region of the AD-fusion triple-helical peptides.

**Fig. 3.**
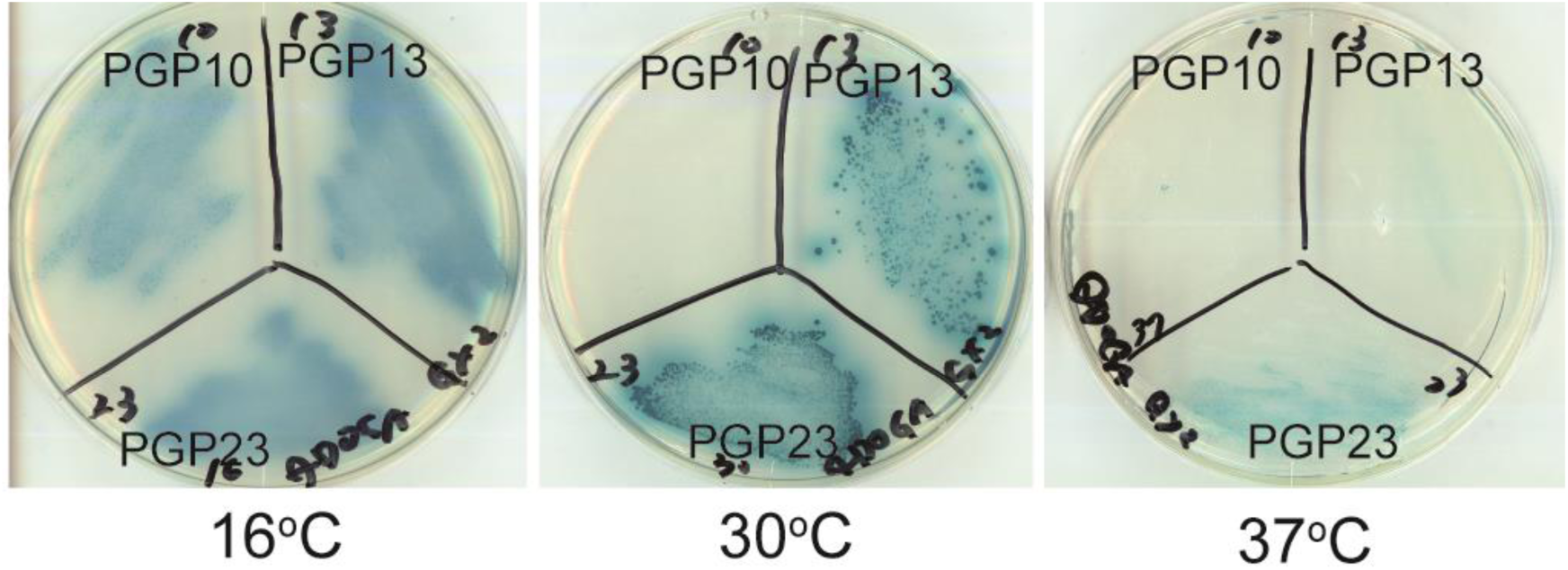
Two-hybrid interaction of HSP47 with (Pro-Gly-Pro)-repeating peptides. PGP10, PGP13, and PGP23 are GAL4-AD-fusion (Pro-Gly-Pro)_10_, (Pro-Gly-Pro)_13_, and (Pro-Gly-Pro)_23_, respectively. They were cultured at 16, 30, or 37°C for 3 days.

### Y2H selection of PEDF-binding peptides from the random peptide library

Two-hybrid screening of the random peptide library was conducted using PEDF as a target protein. PEDF is a collagen-binding serpin that exhibits anti-angiogenic and neurotrophic activities (29–31). It binds to the Lys-Gly-His-Arg-Gly-Phe-Ser-Gly-Leu sequence at colα1(I) 930–938 (15,32). The motif on the triple helix for binding to PEDF was identified as Lys-Gly-Xaa-Arg-Gly-Phe-Yaa-Gly-Leu by Ala-scanning of α1(I)930–938 (15).

The full-length mouse PEDF was expressed as a GAL4-BD-fusion protein in yeast cells and used as bait for the screening. The selection of 1.2 × 10^6^ clones at 25°C yielded three hit clones. Their guest sequences were as follows: Lys-Gly-Cys-Arg-Gly-Leu-His-Gly-Leu (named 1-1), Arg-Gly-Val-Arg-Gly-Phe-Glu-Gly-Cys (named 1-2), and Arg-Gly-His-Arg-Gly-Phe-Leu-Gly-Leu (named 1-3) (Fig. 4A). Although none of these sequences matches those reported in NCBI and UniProt databases, the above three hit clones satisfied the reported structural requirements for binding to PEDF (15): basic amino acids at Yaa1, Arg at Yaa4, and hydrophobic amino acids at Xaa6. Leu at Xaa9, whose replacement by Ala in α1(I)930–938 had led to a partial reduction of the binding affinity to PEDF (15), was also found in the 1-1 and 1-3 sequences.

**Fig. 4.**
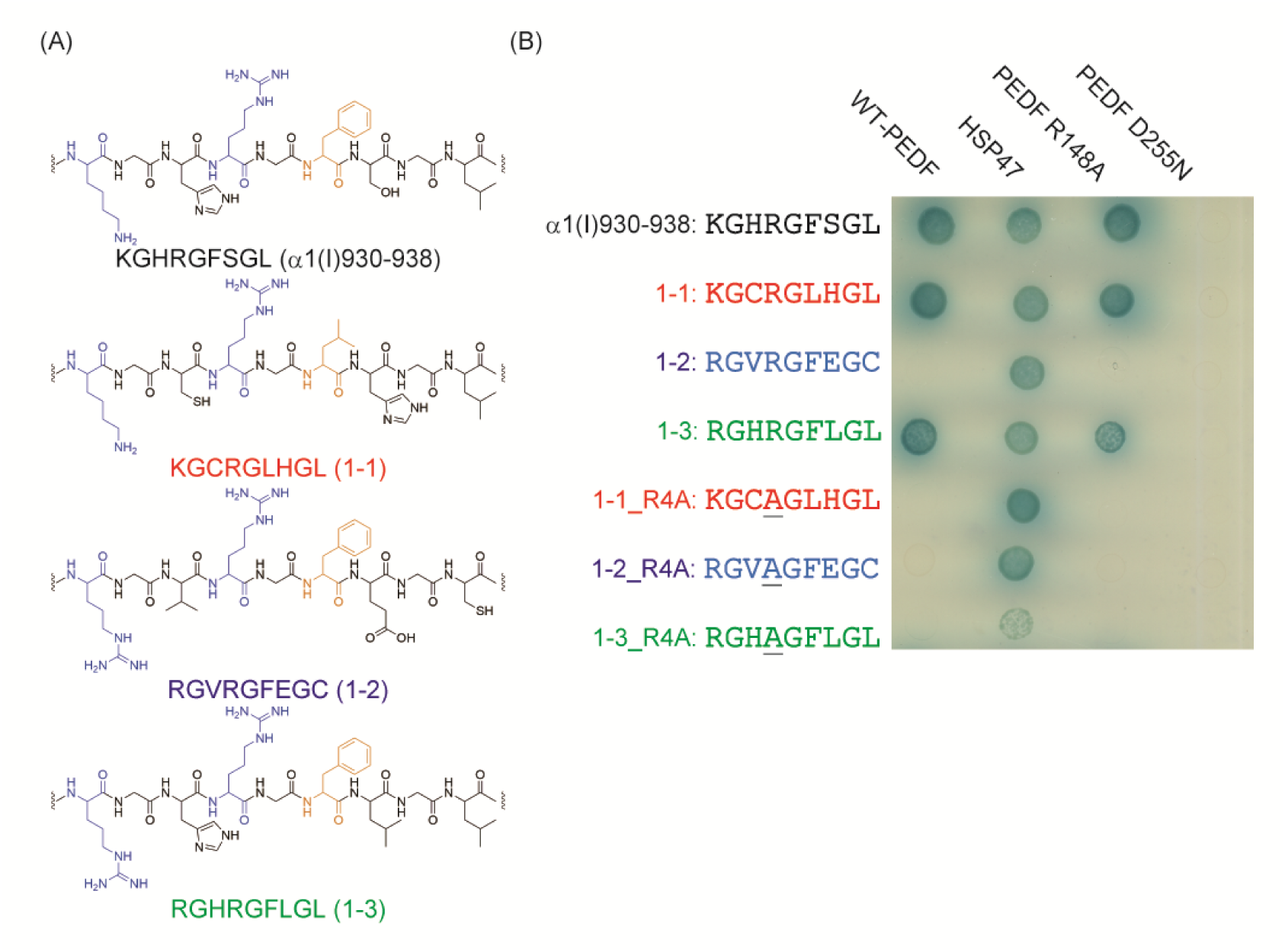
Selection of PEDF-binding triple-helical peptides. (A) The guest sequences of the clones obtained by the selection and native PEDF-binding sequence in type I collagen. (B) The two-hybrid interactions of the AD-fusion peptides with the sequences obtained and their mutants with BD-fusion WT-PEDF, its mutants (R148A and D255N), and HSP47. The cells were cultured at 25°C for 5 days. The sequences of peptides are shown as one-letter codes of amino acids. In the sequences, mutated amino acids are underlined.

To determine whether the peptides with the obtained sequences bind to PEDF in the same manner as the native PEDF-binding sequence, α1(I)930–938, we examined independent two-hybrid interactions of the three peptides and their mutants in which Arg at Yaa4 was substituted with Ala. The peptide with an α1(I)930–938 sequence in which Arg at Yaa4 is substituted with Ala does not bind to PEDF (15). We also examined the binding of the peptides to collagen-binding-null or heparin-binding-null PEDF mutants that we already reported (33). As shown in Fig. 4B, interactions with the AD-fusion peptides with 1-1 and 1-3 sequences, as well as α1(I)930–938, were observed. Meanwhile, the interaction was not observed for the peptide with the 1-2 sequence, probably because Cys at Xaa9 led to weaker interaction with PEDF than Leu (15). The peptide mutants substituted with Ala at Yaa4 (1-1_R4A and 1-3_R4A) did not show interactions with PEDF. In addition, no interactions were observed in the assay using the collagen-binding-null PEDF mutant (PEDF D255N). By contrast, the interactions were maintained using heparin-binding-null PEDF mutant (PEDF R148A) as a bait protein. Besides, the two-hybrid interaction with HSP47 was observed for all peptides. Because we already demonstrated that HSP47 bound to host (Pro-Gly-Pro)-repeating sequences on the triple helix (Fig. 3), all the peptides were strongly suggested to form the triple helix in yeast cells at the assay temperature of 25°C. These results indicated that the two-hybrid selection from the peptide library yielded new PEDF-binding sequences that are recognized in the same manner as those in the native collagen.

### The second-round selection of PEDF-binding peptides from a focused library

Narrowing down the searchable chemical space after acquiring essential structural elements from the screening of a totally random (and hence broad) library is a useful strategy. All three positive clones obtained by the first selection for PEDF-binding peptides contained Arg at the Yaa4 position. Furthermore, the importance of this amino acid residue had been evidenced by the crystal structure of the PEDF-peptide complex (32). We thus further constructed the second library in which the Arg residue was fixed, and the other five residues at the Xaa and Yaa positions were randomized. The guest region of the library encodes the peptide with Yaa1-Gly-Xaa3-Arg4-Gly-Xaa6-Yaa7-Gly-Xaa9. The selection by two-hybrid interaction with PEDF from among 2.1 × 10^6^ clones yielded 52 additional ones, and 12 individual amino acid sequences were obtained as new candidates for PEDF-binding sequences (Table 1). The interaction between the peptides with these amino acid sequences and PEDF was confirmed by two-hybrid assay of the individual clones (Fig. S2). All clones generated blue colonies on the plates. This indicated that the AD-fusion peptides with the sequences obtained from the focused library bind to BD-fusion PEDF.

**Table 1.** Guest sequences of the clones obtained from the focused second library.

| Name | Guest sequence | Frequency |
| --- | --- | --- |
| 2-4 | Arg-Gly-Ala-Arg-Gly-Leu-His-Gly-Leu | 29/52 |
| 2-5 | Arg-Gly-Pro-Arg-Gly-Leu-Thr-Gly-Phe | 8/52 |
| 2-6 | Asn-Gly-Arg-Arg-Gly-Phe-Met-Gly-Met | 3/52 |
| 2-7 | Lys-Gly-Arg-Arg-Gly-Phe-His-Gly-Leu | 2/52 |
| 2-8 | Arg-Gly-Pro-Arg-Gly-Leu-Leu-Gly-Leu | 2/52 |
| 2-9 | Arg-Gly-Phe-Arg-Gly-Leu-Met-Gly-Leu | 1/52 |
| 2-10 | Arg-Gly-Pro-Arg-Gly-Phe-Thr-Gly-Phe | 1/52 |
| 2-11 | Val-Gly-Arg-Arg-Gly-Leu-Ser-Gly-Leu | 1/52 |
| 2-12 | Thr-Gly-Arg-Arg-Gly-Leu-Cys-Gly-Leu | 1/52 |
| 2-13 | Cys-Gly-Val-Arg-Gly-Leu-Thr-Gly-Arg | 1/52 |
| 2-14 | Cys-Gly-Val-Arg-Gly-Leu-Val-Gly-Met | 1/52 |
| 2-15 | His-Gly-Arg-Arg-Gly-Leu-Cys-Gly-Leu | 1/52 |

They are all new amino acid sequences—none of them is found in the NCBI and UniProt databases. Overall, the conventional binding motif, Lys, Arg, or His at Yaa1 and Phe or Leu at Xaa6, was conserved in seven sequences, 2-4, 2-5, 2-7, 2-8, 2-9, 2-10, and 2-15 (15). Meanwhile, the other five sequences, 2-6, 2-11, 2-12, 2-13, and 2-14, did not have a basic amino acid at Yaa1, which suggested that these sequences would interact with PEDF in a different manner from α1(I)930–938.

### Relative PEDF-binding affinity of the chemically synthesized peptides with the sequences obtained from the selections

The amino acid sequences obtained by the two-step yeast two-hybrid selections for PEDF binders were reconstructed as chemically synthesized triple-helical peptides, and their interactions with recombinant PEDF were evaluated. The peptides were designed as host-guest peptides with the sequence of H-Tyr-(Pro-Hyp-Gly)_4_-Pro-[guest sequence]-Hyp-Gly-(Pro-Hyp-Gly)_4_-NH_2_. An N-terminal Tyr residue was introduced to measure the concentration based on the absorbance at 280 nm (34). As a guest sequence, we used the native PEDF-binding sequence and the sequences obtained by the yeast two-hybrid selection, omitting Cys-containing ones to avoid possible oxidative oligomerization. We also prepared **pepfreq**, a peptide whose guest sequence at each position was composed of the amino acid residue most frequently occurring in the two-hybrid selection (Table 2).

**Table 2.** Melting temperature of the triple helix (*T*_m_) and IC_50_ values of the synthetic peptides.

H-Tyr-(Pro-Hyp-Gly)<sub>4</sub>-Pro-[guest sequence]-Hyp-Gly-(Pro-Hyp-Gly)<sub>4</sub>-NH<sub>2</sub>
| Name | Guest sequence | $T_m$<br>(°C) | $IC_{50}^a$<br>(μM) |
| --- | --- | --- | --- |
| **pepα1(I)930-938** | Lys-Gly-His-Arg-Gly-Phe-Ser-Gly-Leu | 37.0 | $0.94 \pm 0.12$ |
| **pep3** | Arg-Gly-His-Arg-Gly-Phe-Leu-Gly-Leu | 43.6 | $4.7 \pm 1.2$ |
| **pep4** | Arg-Gly-Ala-Arg-Gly-Leu-His-Gly-Leu | 47.6 | $1.8 \pm 0.3$ |
| **pep5** | Arg-Gly-Pro-Arg-Gly-Leu-Thr-Gly-Phe | 53.8 | $7.0 \pm 0.8$ |
| **pep6** | Asn-Gly-Arg-Arg-Gly-Phe-Met-Gly-Met | 37.0 | $2.5 \pm 0.3$ |
| **pep7** | Lys-Gly-Arg-Arg-Gly-Phe-His-Gly-Leu | 35.0 | $0.55 \pm 0.66$ |
| **pep8** | Arg-Gly-Pro-Arg-Gly-Leu-Leu-Gly-Leu | 51.0 | >10 |
| **pep9** | Arg-Gly-Phe-Arg-Gly-Leu-Met-Gly-Leu | 50.2 | >10 |
| **pep10** | Arg-Gly-Pro-Arg-Gly-Phe-Thr-Gly-Phe | 57.6 | $0.63 \pm 0.74$ |
| **pep11** | Val-Gly-Arg-Arg-Gly-Leu-Ser-Gly-Leu | 41.8 | $2.8 \pm 0.5$ |
| **pepfreq** | Arg-Gly-Arg-Arg-Gly-Leu-His-Gly-Leu | 44.4 | $1.4 \pm 0.2$ |

<sup>a</sup> $IC_{50}$ values are mean ± S.E.M.

The peptides were synthesized by Fmoc-based solid-phase synthesis and purified by RP-HPLC (Fig. S3 and S4). The conformational state of the synthetic peptides was analyzed by circular dichroism (CD) spectrometry. All of the peptides showed positive signals at approximately 225 nm, indicating that they formed polyproline II helices (35). In addition, the [θ]_225_ signal with increasing temperature cooperatively decreased (Table 2 and Fig. S5). It indicates that the peptides formed triple helices. The peptides were found to maintain the triple-helical structure at 25°C, set in the following PEDF-binding assay, because [θ]_225_ signals at 25°C retained the top positions in sigmoidal curves.

We performed a competitive enzyme-linked immunosorbent assay (ELISA) to evaluate the relative binding affinity of the synthetic peptides to PEDF. In this assay, a competitive decrease in the binding of glutathione *S*-transferase (GST)-PEDF-fusion protein to the type I collagen-coated wells by the addition of the peptides was quantified by immuno-enzymatic colorimetric detection. All the peptides showed concentration-dependent inhibition for PEDF-collagen interaction (Fig. 5). The relative PEDF-binding affinity of the peptides was evaluated using the IC_50_ values derived from the binding inhibitory curves (Table 2 and Fig. 5). Assuming that each trimeric peptide binds to PEDF as 1:1 ratio as well as **pepα1(I)930–938** in the crystal structure (32), **Pep7** exhibited more potent binding affinity than **pepα1(I)930–938** (*p* = 0.0287). **Pep10** and **pepfreq** exhibited binding affinities comparable to that of **pepα1(I)930–938**.

**Fig. 5.**
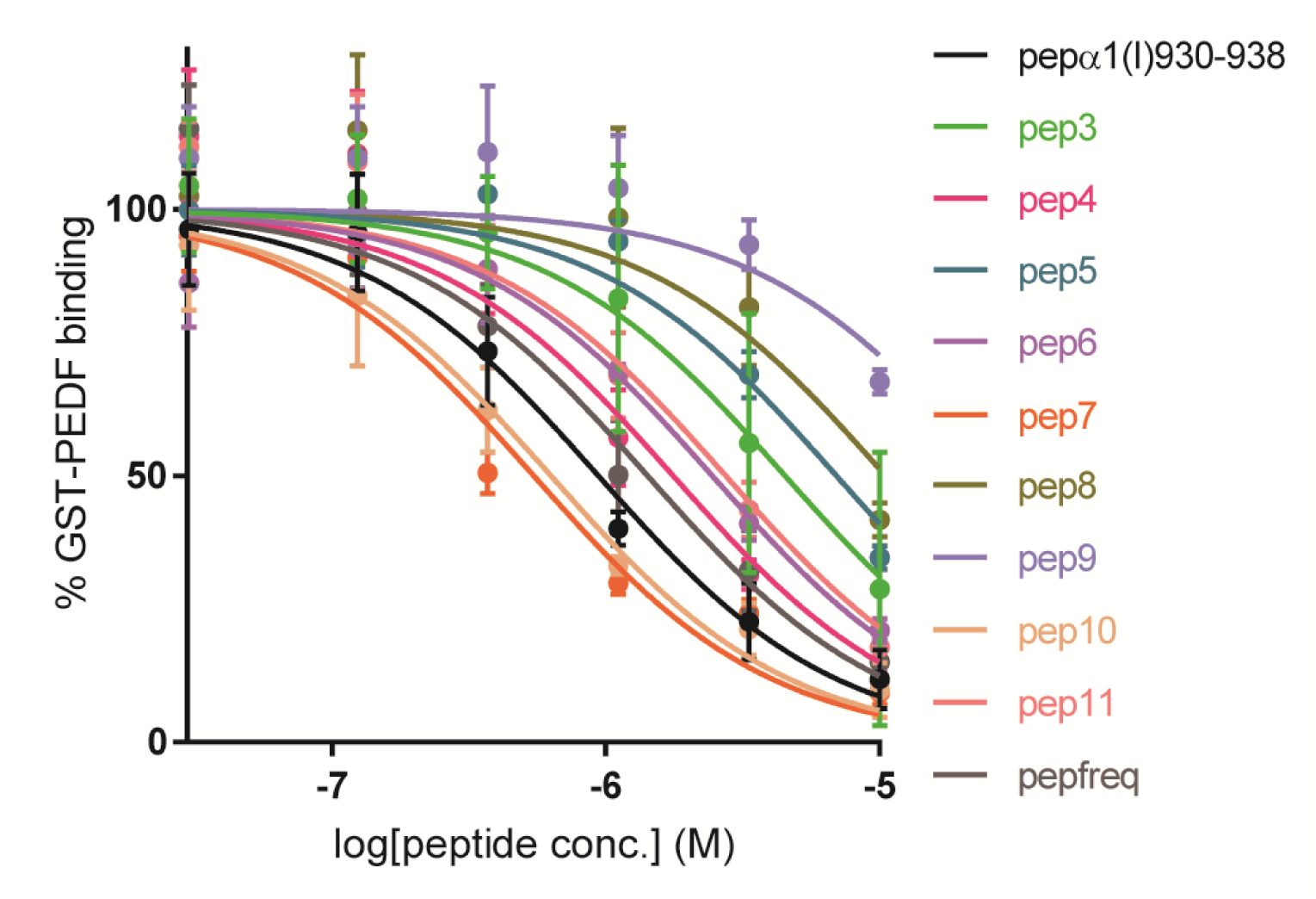
Competitive binding of GST-PEDF to surface-immobilized collagen in the presence of the synthetic peptides. n = 3, mean ± S.D.

### Prediction of a new salt bridge enhances the binding affinity: a docking model simulation

Some of the sequences obtained from the selections, such as 2-6, did not have basic amino acids at Yaa1 but at Xaa3, which does not correspond to the conventional PEDF-binding motif, Lys-Gly-Xaa-Arg-Gly-Phe-Yaa-Gly-Leu. In addition, **pep7** showed higher binding affinity to PEDF than **pepα1(I)930–938**. To investigate the propensity of these peptides to bind to PEDF, we calculated the optimized complex structure of PEDF with **pep6** (Asn1-Gly2-Arg3), **pep10** (Arg1-Gly2-Pro3), and **pep7** (Lys1-Gly-Arg3) from the crystallographic data by molecular modeling. The calculation suggested that Arg3 in **pep6** and Arg1 in **pep10** interacted with the PEDF acidic patch (red color in Fig. 6) that interacts with Lys1 in **pepα1(I)930–938** (Fig. 6A–C). In the calculation of **pep7**, Lys1 pointed toward the acidic patch, like Arg1 in **pep10**. Meanwhile, unlike Arg3 in **pep6**, Arg3 in **pep7** was found to form a new salt bridge with the other acidic amino acids in PEDF (Fig. 6D). This suggests that **pep7** binds to PEDF in a different manner from **pepα1(I)930–938**.

**Fig. 6.**
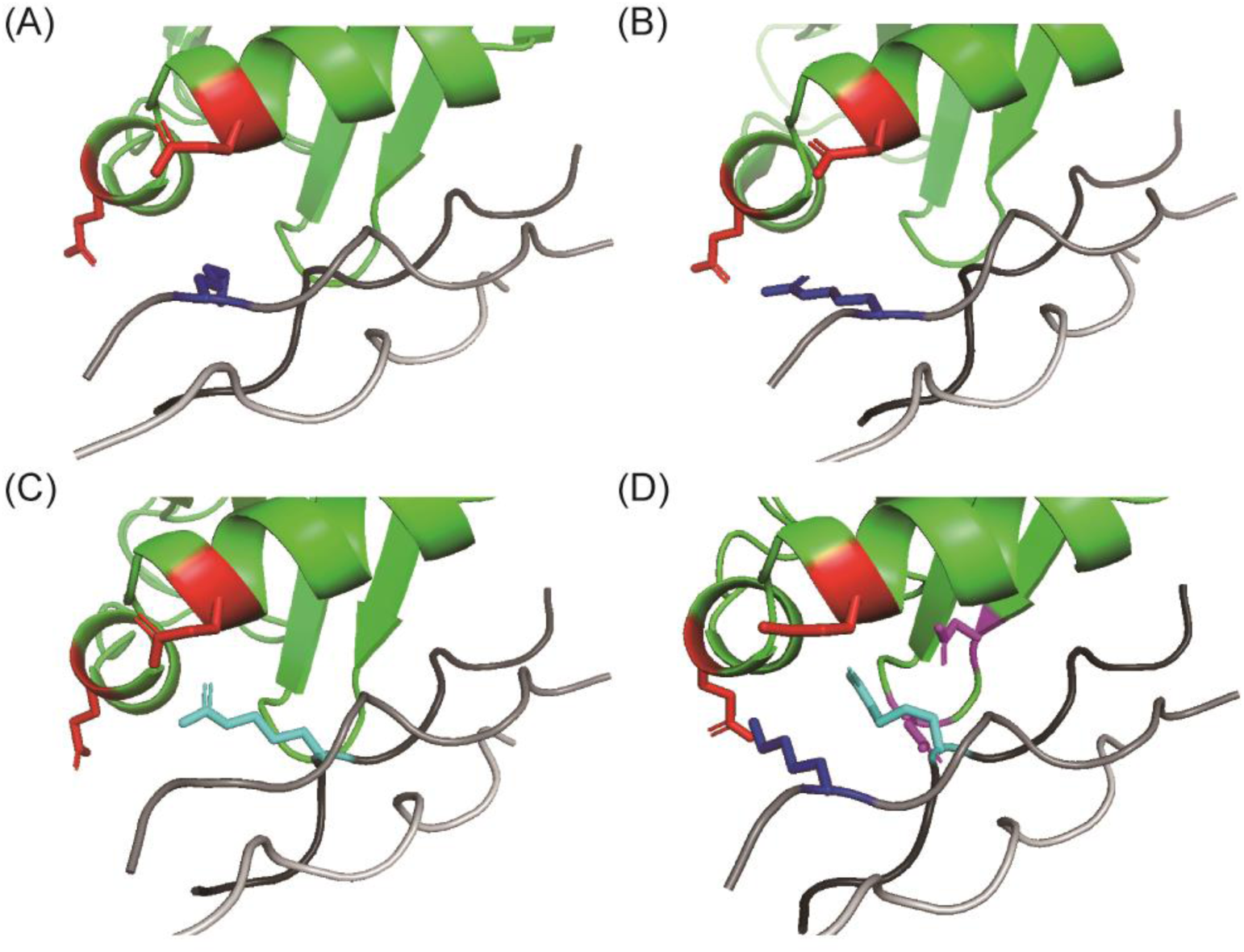
Calculated models of the interactions between PEDF and the peptides obtained [(A) **pepα1(I)930–938**, (B) **pep10**, (C) **pep6**, and (D) **pep7**]. PEDF and the peptides are colored green and gray, respectively. Glu290 and Glu295 in PEDF form the acidic patch with which Lys1 in **pepα1(I)930–938** interacts and are colored red. Asp255 and Asp257 in PEDF are colored magenta. Lys1/Arg1 and Arg3 are colored blue and cyan, respectively. The model was calculated from the reported crystal structure (34, PDB ID: 6LOS).

## Discussion

This work demonstrated the yeast two-hybrid selection of triple-helical peptides that bind to a certain target protein, PEDF, from random combinatorial libraries containing collagen-like Gly-Xaa-Yaa-repeating peptides. The first blind screening without prior knowledge of the binding peptides yielded a few peptides that bind to PEDF within two weeks. The structures of the selected peptides matched the previously identified PEDF-binding amino acid sequence motif as well as the structural requirements revealed by analysis of the co-crystal of the PEDF-triple-helical peptide complex (32). The second round screening of the focused library, in which the essential Arg residue was fixed at the fourth amino acid residue of the randomized portion, resulted in 12 additional peptides. As demonstrated here, the combinatorial use of the completely random first screening and the subsequent second selection from a focused library is a powerful strategy for identifying binding motifs that can be recognized by a POI. By fixing one essential amino acid residue, the coverage of screening theoretically rises 20-fold (in this study, the library coverage increased from 6.3% to 66%). None of the sequences obtained by these two screenings were present in the context of natural collagen α-chains. Our selection system allowed us to obtain 2-7 and 2-10, comparable to or slightly stronger than the potent binding sequence in nature, α1(I)930–938 (Table 2). Considering the coverage of the focused library, 66%, α1(I)930–938 was suggested to be one of the optimized sequences for potent binding to PEDF by natural selection.

Screening from the random triple-helical library yielded several sequences that do not correspond to the conventional motif for binding to PEDF. The docking simulation suggests that Arg3 of **pep6** points toward the acidic patch, like Arg1 of **pepα1(I)930–938**. Meanwhile, although Arg1 of **pep7** retains the interaction with the acidic patch, Arg3 of **pep7** additionally interacts with a different PEDF acidic patch with which His3 of **pepα1(I)930–938** does not interact (Fig. 6). **Pep7** exhibited more potent binding to PEDF than **pepα1(I)930–938**, probably because of this additional interaction. This analysis suggests that the yeast two-hybrid system is useful for finding non-natural motifs for binding to collagen-binding proteins.

The procedure of the screening based on the standard yeast two-hybrid technique is simple and easy to access without any purification or immobilization of proteins or peptides. Screening of 10^6^ clones out of the theoretical diversity of 20^6^ can be conducted in a single-round selection. The system relies on the self-trimerizing propensity of the peptides consisting of collagenous Gly-Xaa-Yaa repeats, and on following oligomeric interaction with the target protein in the yeast nuclei. By taking advantage of the host-guest design that stabilizes the triple-helical conformation, we achieved an increase in the population of triple-helical molecules in the library. The melting temperature of **pep7** (*T*_m_ = 35.0°C) was lower than that of **pepα1(I)930–938** (*T*_m_ = 37.0°C), whose guest sequence is known as the most unstable region of type I collagen (19). This indicates that the peptide design enabled the library to contain even amino acid sequences that are too unstable to form a triple helix in the context of natural collagen. As the yeast two-hybrid system allows us to conduct screening at temperatures ranging from 16°C to 37°C (Fig. 3), the population of the triple-helical molecules in the library can be expected to increase by lowering the screening temperature. Even if the selected sequences cannot form triple helices by themselves at the body temperature, it is possible to present and use them as triple helix peptides. Various peptide designs have been developed to enhance the thermal stability of the triple helix, such as host sequence optimization and chemical tethering of the three chains, which are readily accessible to researchers (3,36,37).

The two-hybrid interaction with HSP47 was shown to be useful in confirming the triple-helical conformation of the peptides expressed in the yeast clones (Figs. 3 and 4B). It is by virtue of the conformational-specific recognition of the collagen-like peptides by the chaperone. This unique property would also be utilized for the quality control of the triple-helical peptide library constructed in the yeast cells by performing pre-screening by HSP47.

Collagen Toolkits have contributed to identifying the amino acid sequences on collagen triple helices responsible for the protein interactions (5–12). From these previous studies, we realized that there is a large overlap in recognition among different collagen-binding proteins. As the most typical case, VWF, DDR, SPARC, and aegyptin, a mosquito salivary gland protein, were found to share the same binding site in native collagen (Table S1) (10–13,38). One reason for the peculiar interactions between collagen and proteins is the low diversity of amino acid sequences displayed on the triple helices of native collagen. As shown in Fig. 2, 226 Xaa-Yaa combinations appeared in the fibril-forming collagen out of 400 (= 20 × 20) possible combinations, and they tended not to include triple-helix-destabilizing Xaa-Yaa combinations. The use of the more diverse combinatorial peptide libraries unrelated to the consequences of the molecular evolution of collagen raised expectations of obtaining selective binders for individual collagen-binding proteins. Such selective binders would be useful tools for resolving the complex assortment of interactions between proteins and collagen and hence in elucidating collagen’s physiological regulatory functions. On the other hand, a limitation of the system is that Pro residues in the peptide are not hydroxylated to form Hyp residues because of the absence of the collagen prolyl 4-hydroxylase in yeast cells.

The triple-helical peptides that specifically bind to bioactive proteins have the potential to be used as lead compounds of drug development. Because chemically synthesized triple-helical peptides show remarkable stability in living animals by resisting attacks by proteases (39,40), the triple helix is a promising scaffold for peptide drugs. Alternatively, the triple-helical peptides with protein-binding sequences obtained by the screening can be used in artificial collagen-like polymer systems to make novel functional biomaterials (41–44). As an additional reward, targets are not limited to the collagen-binding proteins; any proteins could serve as bait for the selection.

## Experimental procedures

### Construction of the plasmids encoding GAL4-BD-fused proteins

A gene fragment encoding full-length mature mouse PEDF and HSP47 with *EcoR*I and *Sal*I sites was obtained by PCR amplification, as described in the literature (15,45). The DNA fragments and a pGBKT7 vector were ligated after digestion with *EcoR*I and *Sal*I and transformed into *E. coli* HST08 competent cells (TaKaRa, Shiga, Japan). To prepare plasmids encoding PEDF R148A and D255N mutants, inverse PCR was performed with the above GAL4-BD-fused PEDF in a pGBKT7 vector as a template by Primestar MAX (TaKaRa). The PCR product was ligated with Gibson assembly reagents (New England Biolabs, Hertfordshire, UK), followed by transformation into HST08 competent cells. Used primers are listed in Table S2.

### Construction of plasmids encoding GAL4-AD-fused peptides

To prepare a plasmid encoding GAL4-AD-fused peptide with α1(I)930–938, (Pro-Gly-Pro)_10_-Lys-Gly-His-Arg-Gly-Phe-Ser-Gly-Leu-(Pro-Gly-Pro)_10_ attached with *EcoR*I and *BamH*I sites in a pUCFa vector was synthesized by FASMAC (Kanagawa, Japan). The DNA fragment and a pGADT7 AD vector were ligated with *EcoR*I and *BamH*I after digestion and transformed into HST08 competent cells. The plasmid, named α1(I)930–938/pGADT7, was extracted from the cells. The sequence map of α1(I)930–938/pGADT7 is shown in Fig. S6.

To prepare plasmids encoding GAL4-AD-fused PGP10, PGP13, PGP23, and peptides with 1-1_R4A, 1-2_R4A, and 1-3_R4A sequences, α1(I)930–938/pGADT7 was transformed into *E. coli* HST04 (dam-/dcm-) competent cells (TaKaRa) to avoid DNA methylation digesting the *Apa*I site. The plasmid was extracted from the cells, followed by digestion with *Apa*I and *Xma*I. As preparation for the insertion of DNA fragments, we annealed a single-stranded DNA (ssDNA) encoding corresponding peptide sequences attached with *Apa*I and *Xma*I cut sites with an antisense ssDNA at 95°C for 5 min (Table S3). After cooling at room temperature overnight, the annealed DNA was ligated and transformed into HST08 competent cells.

To obtain plasmids from yeast cells, yeast clones were lysed to extract DNA using Zymoprep Yeast Plasmid Miniprep I kit (Zymo Research, Irvine, CA, USA), in accordance with the manufacturer’s protocol. The DNA was transformed into HST08 competent cells and inoculated on LB plates with ampicillin to select prey constructs.

### Transformation of DNA into yeast cells, mating, and detection of Y2H interactions

The bait and prey plasmids were transformed into *S. cerevisiae* Y2H gold (*MATa, trp1-901, leu2-3, 112, ura3-52, his3-200, gal4Δ, gal80Δ, LYS2::GAL1_UAS_-Gal1_TATA_-His3, GAL2_UAS_-Gal2_TATA_-Ade2, URA3::MEL1_UAS_-Mel1_TATA_, AUR1-C MEL1*; TaKaRa) and Y187 strains (*MATα, ura3-52, his3-200, ade2-101, trp1-901, leu2-3, 112, gal4Δ, gal80Δ, met-, URA3::GAL1_UAS_-Gal1_TATA_-LacZ, MEL1*; TaKaRa), in accordance with the manufacturer’s protocol. They were incubated on SD/-Trp and SD/-Leu plates, respectively, at 30°C for 4–5 days for the selection of transformed clones.

Diploids were prepared by mixing them on yeast peptone dextrose adenine (YPDA) plates. After incubation at 30°C overnight, they were spread on minimal-medium double-dropout (DDO; SD/-Leu-Trp) plates and incubated at 30°C for 2–3 days for the selection of diploids.

Y2H interaction was detected by incubation of the generated diploids on minimal-medium quadruple-dropout (QDO; SD/-Leu-Trp-His-Ade) plates with 40 μg/ml X-α-Gal (FUJIFILM Wako Pure Chemical, Osaka, Japan) and 200 ng/ml aureobasidin A (TaKaRa) (QDOGA) plates and incubated for 3–5 days. Positivity for interaction was recorded upon the generation of yeast colonies that appeared blue.

### Preparation of a random triple-helical library and selection of protein-binding peptides

First, 10 μM sense DNA encoding Yaa1-Gly-Xaa3-Yaa4-Gly-Xaa6-Yaa7-Gly-Xaa9 (C NNK GGT NNK GGA NNK GGT NNK C) and 10 μM corresponding antisense DNA (CC GGG MNN ACC MNN CCT MNN ACC MNN GGG CC) were annealed in the presence of 300 mM NaCl at 95°C for 5 min (N is an equimolar mixture of A/T/G/C, K is an equimolar mixture of T/G, and M is an equimolar mixture of A/C). The sample was then cooled down to room temperature for 1 h. The annealed DNA was ligated with pGADT7 after digestion with *Apa*I and *Xma*I by T4 DNA ligase (Takara) at 16°C for 1 h (insert:vector molar ratio is 10:1), and transformed into JetGiga *E. coli* cells (Funakoshi, Tokyo, Japan), in accordance with the manufacturer’s protocol. DNA was extracted from the colonies growing on LB plates with ampicillin using Midiprep kit (Sigma-Aldrich, St. Louis, MO, USA) and transformed into the Y187 strain, in accordance with the manufacturer’s protocol. Colonies growing on SD/-Leu plates were collected with 25% glycerol YPDA, divided into tubes (5.6 × 10^6^ yeast cells/tube), and stored at −80°C until use.

For selection, the protein-expressing Y2H gold strain was cultured to 0.5–0.6 OD_600_ in 50 mL of SD/-Trp broth at 30°C. This culture was mixed with one tube of the library stocks and filtrated through a nylon membrane filter (0.45 μm pore; GE Healthcare Life Science, Pittsburgh, PA, USA). After incubation of the membrane on a YPDA plate at 30°C overnight, yeast was collected with saline and inoculated on QDO plates with 200 ng/ml aureobasidin A. Positive clones were determined by the appearance of colonies with a blue color on QDOGA plates after 5 days of incubation at 25°C. Their guest sequences were identified by analyzing the sequence of PCR products by colony PCR.

### Investigation of Xaa-Yaa combination of the peptides in the library constructed in yeast cells by NGS analysis

Y187 strain of the library was suspended in yeast lysis buffer [50 mM Tris/HCl (pH 8.0), 150 mM NaCl, 5 mM EDTA, 1% NP-40, and 1% SDS] and the same volume of phenol/chloroform/isoamyl alcohol (25:24:1, pH 7.9) was added, followed by vortexing with glass beads for 1 min. After centrifugation at 20,600 × *g* for 5 min, the upper layer was recovered. DNA plasmids were purified from the solution by precipitation with ethanol. PCR products for amplicon sequencing were amplified with the purified DNA and primers attached to adaptor sequences (Table S4). Next-generation sequencing analysis was performed by Macrogen (Kyoto, Japan) with the PCR products. We extracted 27 bases with a nucleotide sequence of NNN GGT NNN GGA NNN GGT NNN flanked by *Apa*I and *Xma*I sites without a stop codon from NGS data (54,848 reads, Dataset_S01). The nucleotide sequences were translated according to the codon table to obtain guest amino acid sequences. Each of the Gly2-Xaa3-Yaa4 and Gly5-Xaa6-Yaa7 triplets was categorized into four groups according to the effect on the thermostability of the triple helix, as previously reported (19).

### Peptide synthesis and characterization

The peptide chain was manually constructed using the 9-fluorenylmethyloxycarbonyl (Fmoc)-based solid-phase method on Rink amide resin (Merck, Darmstadt, Germany). In each cycle, Fmoc-amino acids (5 equivalents) were reacted in the presence of *N,N’*-diisopropylcarbodiimide (5 equivalents) and 1-hydroxybenzotriazole (HOBt) (5 equivalents) in *N,N*-dimethylformamide (DMF). Fmoc deprotection was performed with 20% (v/v) piperidine in DMF for 20 min. The protected peptide resin was treated with trifluoroacetic acid (TFA)/H_2_O/*m*-cresol/thioanisole/triisopropylsilane (82.5/5/5/5/2.5, v/v) for 4 h at room temperature. The crude products were purified by RP-HPLC on a Cosmosil 5C18-AR-II column (8.0 × 250 mm; Nacalai Tesque, Kyoto, Japan) with CH_3_CN in water, both containing 0.05% (v/v) TFA, to afford the expected peptides. The compounds were analyzed using RP-HPLC on a Cosmosil 5C18-AR-II (4.6 × 250 mm; Nacalai Tesque) with 10%−40% CH_3_CN in 0.05% TFA over 30 min at 60°C and absorbance was measured at 220 nm. Mass spectrometric analysis was performed with a Bruker Autoflex III matrix-assisted laser desorption ionization-time-of-flight (MALDI) mass spectrometer (Bruker Daltonics, Leipzig, Germany) or a Triple TOF 4600 (AB SCIEX, Foster City, CA, USA) electrospray ionization mass spectrometer MS. α-Cyano-4-hydroxycinnamic acid was used as a matrix in the MALDI analysis.

### CD spectrometry of the synthetic peptides

CD spectra were recorded on a J-820 CD spectropolarimeter (JASCO, Tokyo, Japan) equipped with a Peltier thermal controller using a 0.5-mm path-length quartz cuvette and connected to a data station for signal averaging. The peptides in PBS were heated at 95°C for 5 min, incubated at room temperature for 10 min, and stored at 4°C overnight to allow formation of the triple-helical structure. Data were obtained using continuous wavelength scans from 200 to 260 nm. The spectra are reported in terms of ellipticity units per mole of amino acid residue [θ]_mrw_. Triple-helix thermostability was monitored by observation of each peptide [θ]_225_ value with increasing temperature from 4 to 85°C at 18°C per hour. The obtained molar ellipticity was differentiated to calculate the slopes at each temperature, and the temperature at which the minimum value was obtained was taken as the melting temperature of the triple helix (*T*_m_).

### Competitive ELISA

GST-fused PEDF was prepared in accordance with a previous report (Fig. S7) (15). The synthetic peptides were annealed by the above method before being used in the assay. Wells were coated with 10 μg/ml collagen I (Koken, Tokyo, Japan) overnight. After coating, wells were blocked with 0.5% skim milk in ELISA buffer [20 mM HEPES-Na (pH 7.4), 100 mM NaCl, 0.005% Tween 20]. The annealed peptides were preincubated with 3 nM GST-PEDF to be tested in the ELISA buffer for 10 min at room temperature. The mixtures were added to wells and incubated for 90 min at 4°C. After washing with ELISA buffer, horseradish peroxidase (HRP)-conjugated anti-GST antibody (diluted 1:3000; GE Healthcare) was added and incubated for 30 min at 4°C. After washing, 0.5 mg/ml 2,2′-azino-bis(3-ethylbenzothiazoline-6-sulfonic acid) diammonium salt (FUJIFILM Wako Pure Chemical) in citrate-phosphate buffer [0.1 M citrate and 0.2 M Na_2_HPO_4_ (pH 5.0)] was added to each well. After 30 min, the absorbance at 405 nm was measured on a Vient XS multiwell plate reader (DS Pharma, Osaka, Japan). Sigmoidal curves were drawn using GraphPad Prism software version 7.04 (GraphPad Software, San Diego, CA, USA) to fit the curves with nonlinear regression. The *p*-value was calculated from an F-test.

### Molecular modeling

The binding modes of **pepα1(I)930–938, pep6**, **pep7**, and **pep10** were evaluated using molecular modeling methods by substituting the residues of CP211 in the crystal structure of PEDF-CP211 complex (PDB code: 6LOS) with the residues corresponding to each peptide. The resulting models were energy-minimized using GROMACS 2020 software package by the quasi-Newton method until the energy of the system converged to 100 kJ mol^−1^ nm^−1^ (46). The energy calculation was performed with the AMBER99SB*-ILDN energy terms under the NPT ensemble and periodic boundary conditions. Simple point-charge water molecules were chosen for the water model with several counterions added to attain an electrically neutral system.

### Database search

To search for the obtained sequences in collagen, a BLAST search was performed using the obtained sequences as queries in NCBI (https://blast.ncbi.nlm.nih.gov/Blast.cgi) and UniProt (https://www.uniprot.org/blast) databases.

## Data Availability

All data is contained within the manuscript.

## Acknowledgments

We thank Edanz (https://jp.edanz.com/ac) for editing a draft of this manuscript. The work was supported by JSPS KAKENHI Grant Number 20K05756, Waseda University Grant for Encouragement of Scientists, and JST SPRING, Grant Number JPMJSP2128.

## Notes

### Competing Interest Statement

The authors have declared no competing interest.

### Summary of Updates

The revised submission updated the clarity and enhanced the significance of our study to obtain the triple-helical ligands from the combinatorial random peptide libraries by the yeast two-hybrid system. Here are the detailed updates of our manuscript. 1. We added a better discussion of recognizing the HSP47 bound (Pro-Pro-Gly)- repeating sequences in their host region as the triple-helix structure at the assay temperature of 25oC. (page 11, lines 1-4). 2. We changed the writing to more comprehensive, describing the concentration measurement by spectrophotometry. (page 12, lines 11-12). 3. To show a comparable binding affinity of the two synthetic peptides with a potent binding sequence in nature, α1(I)930-938, we added a deeper discussion based on the coverage of the focused library 66%. (page 15, lines 15-18). 4. We additionally included more information and references to the impact on the biological relevance of the screened peptides. (page 16, lines 16-20). 5. The more discussion part of the pre-screening with HSP47 was added to have a better understanding of the unique property, the conformational-specific recognition of the HSP47 chaperone, which can be applied to the assurance of the triple-helical peptide libraries constructed in the yeast cells. (page 16, lines 22-23; page 17, lines 1-2). 6. We enlarged the figures of MS data to have a high resolution. (figure S4). 7. Figure 2A was replaced with the correct one. 8. We moved figure 5 to figure S2 in the supporting information. 9. The experimental procedures section was moved after the discussion section. 10. The data availability section was placed after the experimental procedures section.

