## Supporting Information for "A yeast two-hybrid system to obtain triple-helical ligands from combinatorial random peptide libraries"

Table S1. Collagen-binding proteins and the identified binding sequences in human collagen

| Protein | Collagen-binding domain | Binding sequence[a,b] | Ref. |
|---|---|---|---|
| PEDF | single-domain protein | KGHRGFSGL [α1(I)(87–95)/α1(I)(930–938)] | 1 |
| integrin α1β1 | αI domain | GLOGEN [α1(II)(121–126)] [c] | 2 |
| integrin α2β1 | αI domain | GFOGER [α1(I)(502–507)/α1(III)(322–327)] [c] | 3 |
| integrin α10β1 | αI domain | GLOGEN [α1(II)(121–126)] [c] | 4 |
| integrin α11β1 | αI domain | GFOGER [α1(I)(502–507)/α1(III)(322–327)] [c] | 5 |
| VWF | A3 domain | RGQOGVMGF [α1(III)(405–413)] | 6 |
| SPARC | EC domain | GVMGFO [α1(III)(409–414)] | 7 |
| DDR1/2, | DS domain | RGQOGVMGFO [α1(III)(405–414)] | 8,9 |
| aegyptin | -[d] | RGQOGVMGF [α1(III)(405–413)] | 10 |
| MMP-1 | cat domain | OGPQGLAGQR [α1(II)(771–780)] | 11 |
| MMP-3 | cat domain | GAAGFOGAR [α1(III)(533–541)] | 12 |

[a] Text in parentheses indicates the locations in collagen

[b] "O" means 4-hydroxy-L-proline

[c] The most potent sequence in collagen

[d] Not reported

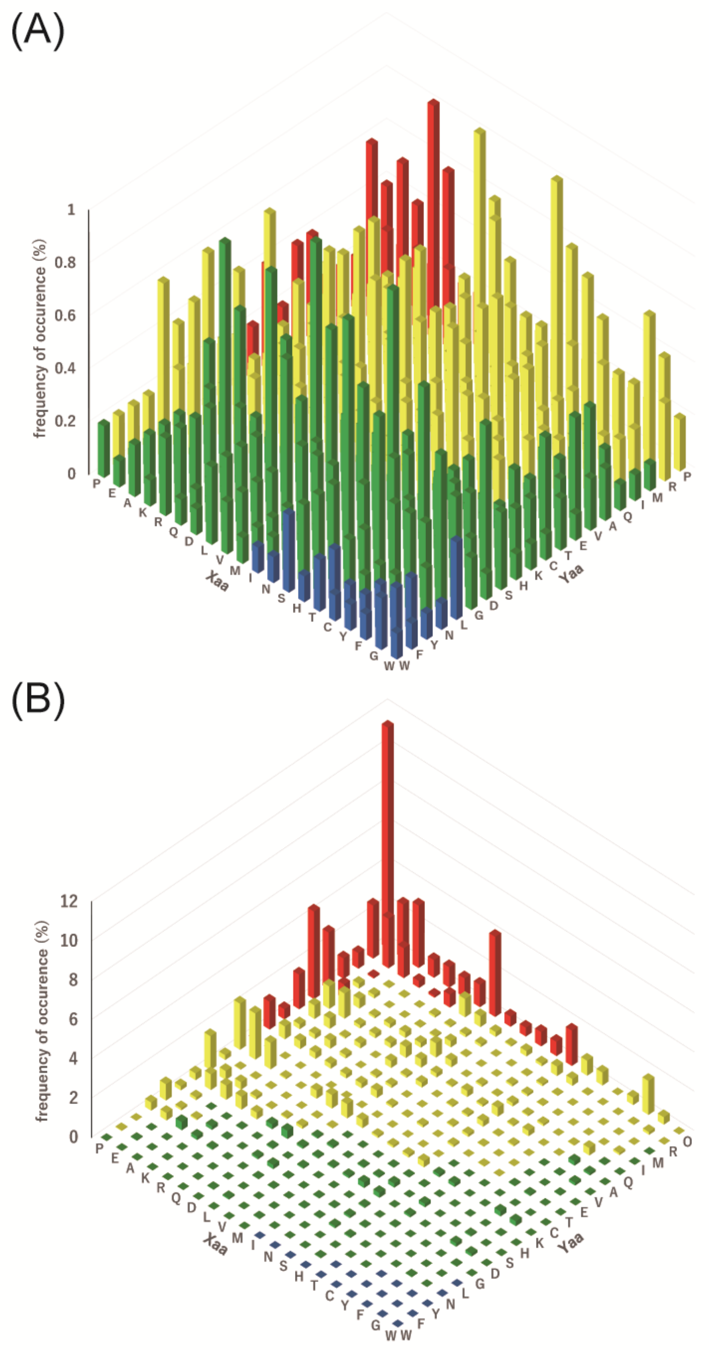


Fig. S1. Distribution of Gly-Xaa-Yaa triplets. (A) Theoretical distribution of triplets in which Xaa and Yaa are encoded by NNK codons. (B) A reduced view of Gly-Xaa-Yaa triplet distribution of human fibrillar collagen. Regarding the change from Fig. 2B to this figure, the maximum value of the Z-axis changes from 1% to 12%.

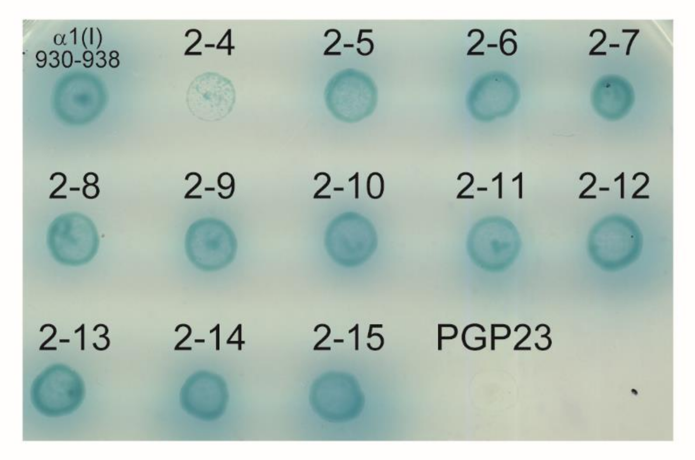


Fig. S2. Interaction of the AD-fusion peptides with the sequences obtained from the second selection with BD-fusion PEDF in yeast cells. PGP23 is GAL4-AD-fusion (PGP)23. The cells were cultured at 25°C for 5 days.

**pepα1(I)930-938**

H-Tyr-(Pro-Hyp-Gly)$_4$-Pro-Lys-Gly-His-Arg-Gly-Phe-Ser-Gly-Leu-Hyp-Gly-(Pro-Hyp-Gly)$_4$-Pro-NH$_2$

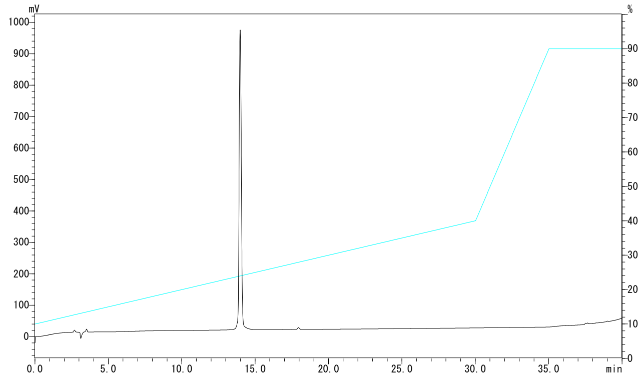


T$_R$: 14.0 min

**pep3**

H-Tyr-(Pro-Hyp-Gly)$_4$-Pro-Arg-Gly-His-Arg-Gly-Phe-Leu-Gly-Leu-Hyp-Gly-(Pro-Hyp-Gly)$_4$-Pro-NH$_2$

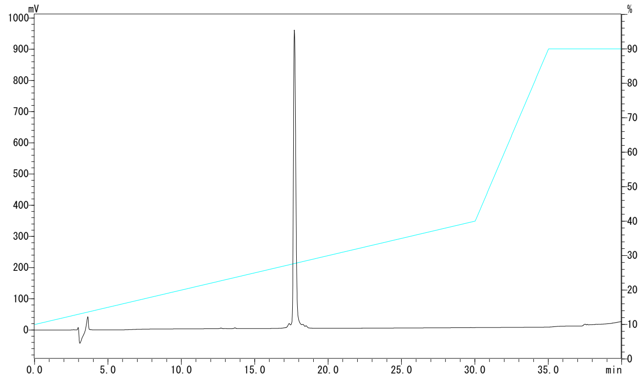


T$_R$: 17.7 min

**pep4**

H-Tyr-(Pro-Hyp-Gly)$_4$-Pro-Arg-Gly-Ala-Arg-Gly-Leu-His-Gly-Leu-Hyp-Gly-(Pro-Hyp-Gly)$_4$-Pro-NH$_2$

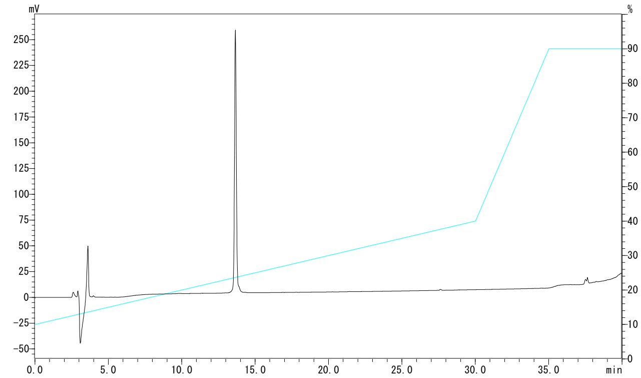


T$_R$: 13.7 min

Fig. S3. HPLC profiles of the synthetic peptides.

**pep5**

H-Tyr-(Pro-Hyp-Gly)$_4$-Pro-Arg-Gly-Pro-Arg-Gly-Leu-Thr-Gly-Phe-Hyp-Gly-(Pro-Hyp-Gly)$_4$-Pro-NH$_2$

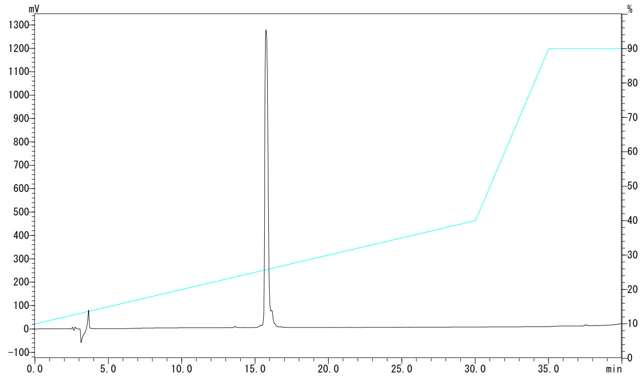


$T_R$: 15.7 min

**pep6**

H-Tyr-(Pro-Hyp-Gly)$_4$-Pro-Asn-Gly-Arg-Arg-Gly-Phe-Met-Gly-Met-Hyp-Gly-(Pro-Hyp-Gly)$_4$-Pro-NH$_2$

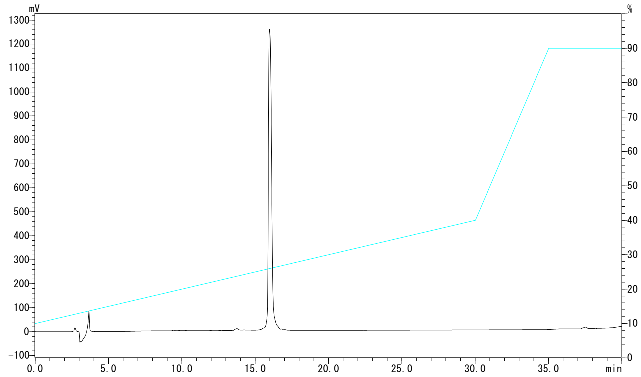


$T_R$: 16.0 min

**pep7**

H-Tyr-(Pro-Hyp-Gly)$_4$-Pro-Lys-Gly-Arg-Arg-Gly-Phe-His-Gly-Leu-Hyp-Gly-(Pro-Hyp-Gly)$_4$-Pro-NH$_2$

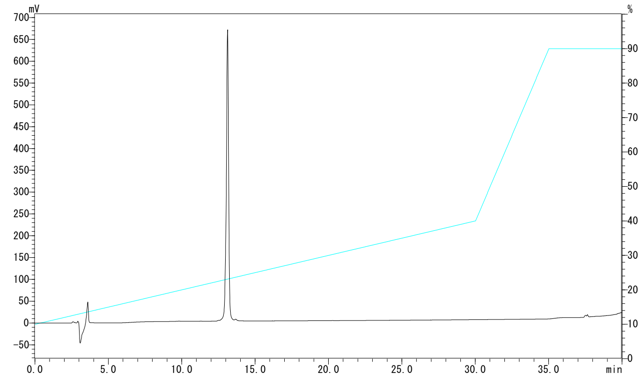


$T_R$: 13.1 min

Fig. S3. (*Continued*).

**pep8**

H-Tyr-(Pro-Hyp-Gly)$_4$-Pro-Arg-Gly-Pro-Arg-Gly-Leu-Leu-Gly-Leu-Hyp-Gly-(Pro-Hyp-Gly)$_4$-Pro-NH$_2$

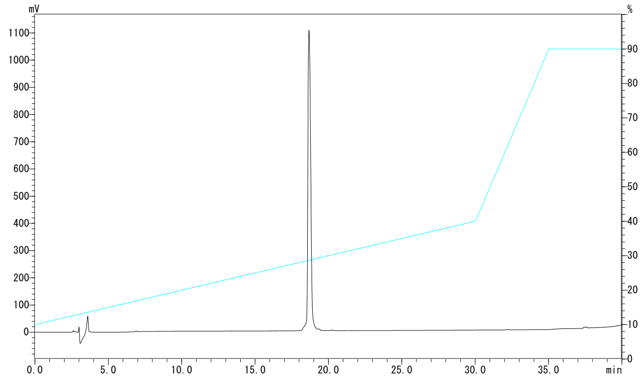


T$_R$: 18.7 min

**pep9**

H-Tyr-(Pro-Hyp-Gly)$_4$-Pro-Arg-Gly-Phe-Arg-Gly-Leu-Met-Gly-Leu-Hyp-Gly-(Pro-Hyp-Gly)$_4$-Pro-NH$_2$

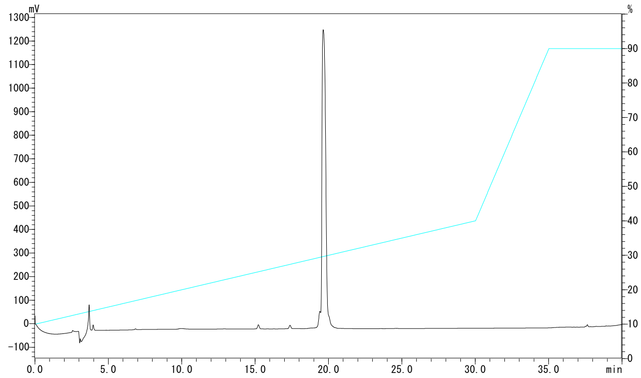


T$_R$: 19.6 min

**pep10**

H-Tyr-(Pro-Hyp-Gly)$_4$-Pro-Arg-Gly-Pro-Arg-Gly-Phe-Thr-Gly-Phe-Hyp-Gly-(Pro-Hyp-Gly)$_4$-Pro-NH$_2$

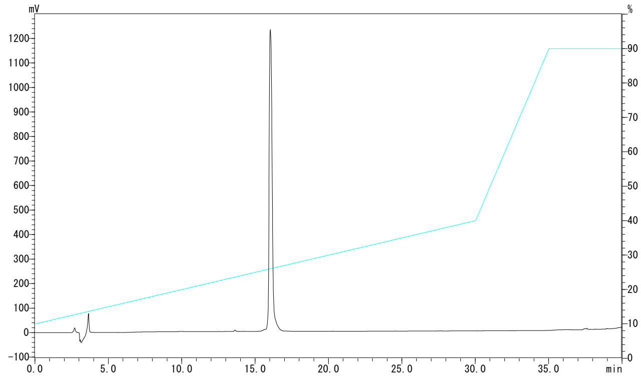


T$_R$: 16.0 min

Fig. S3. (*Continued*).

**pep11**

H-Tyr-(Pro-Hyp-Gly)$_4$-Pro-Val-Gly-Arg-Arg-Gly-Leu-Ser-Gly-Leu-Hyp-Gly-(Pro-Hyp-Gly)$_4$-Pro-NH$_2$

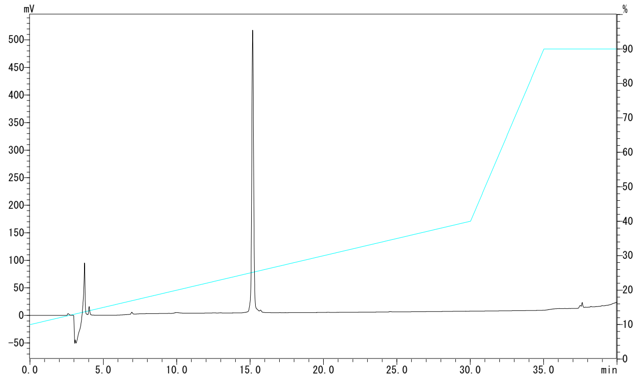


$T_R$: 15.2 min

**pepfreq**

H-Tyr-(Pro-Hyp-Gly)$_4$-Pro-Arg-Gly-Arg-Arg-Gly-Leu-His-Gly-Leu-Hyp-Gly-(Pro-Hyp-Gly)$_4$-Pro-NH$_2$

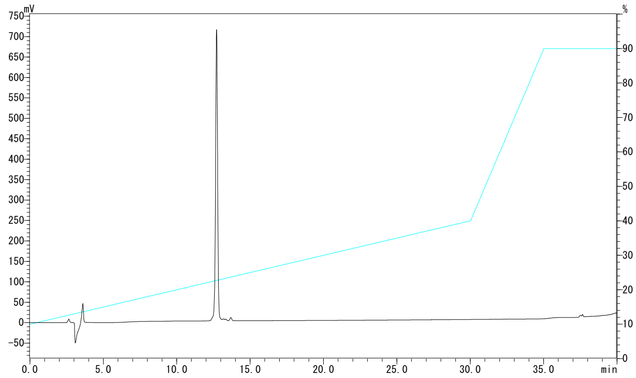


$T_R$: 12.7 min

Fig. S3. (*Continued*).

**pepα1(I)930-938**

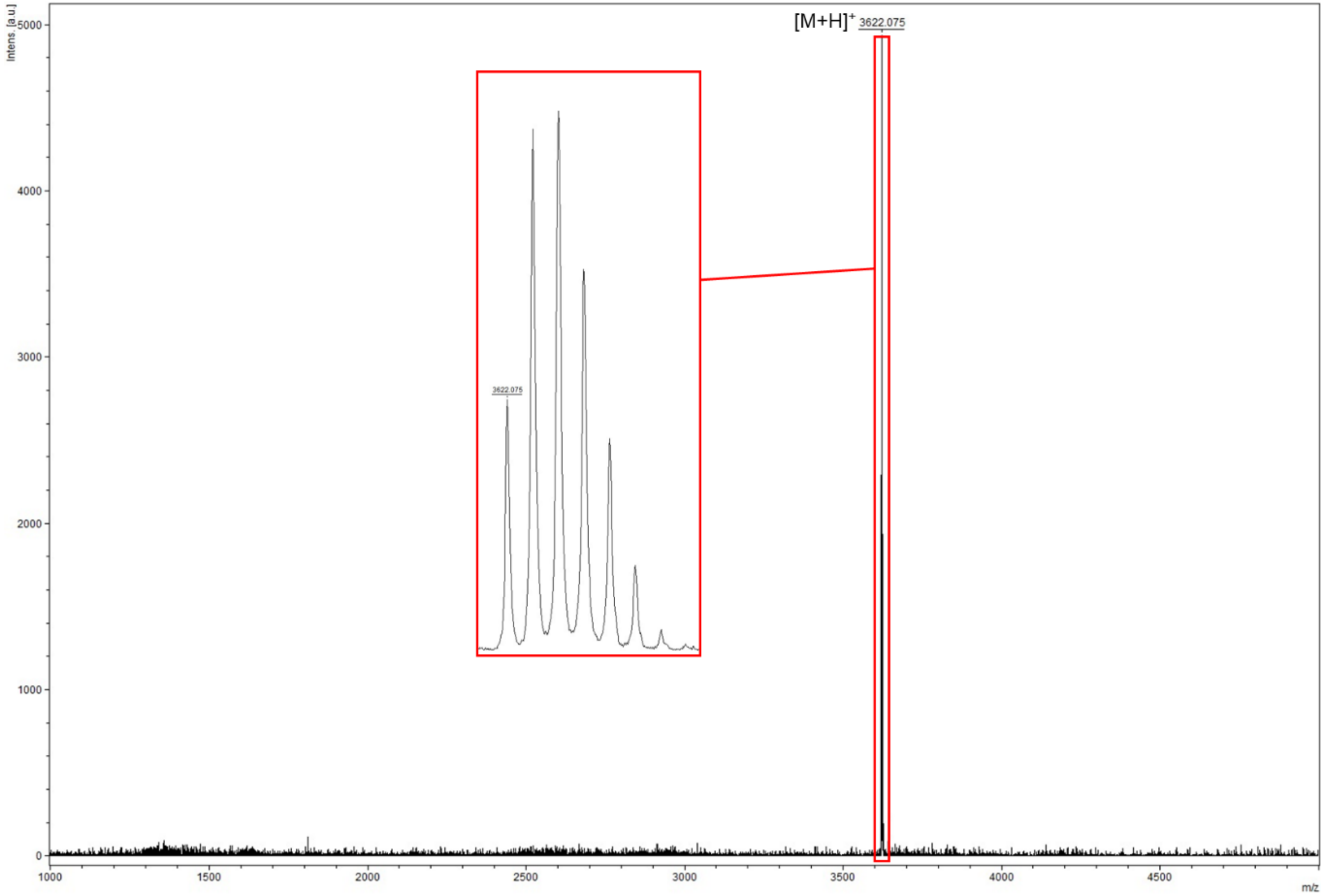


calcd. MS [$M_m$ + H] $^+$: 3621.75            found: 3622.08

Fig. S4. MS charts of the synthetic peptides.

**pep3**

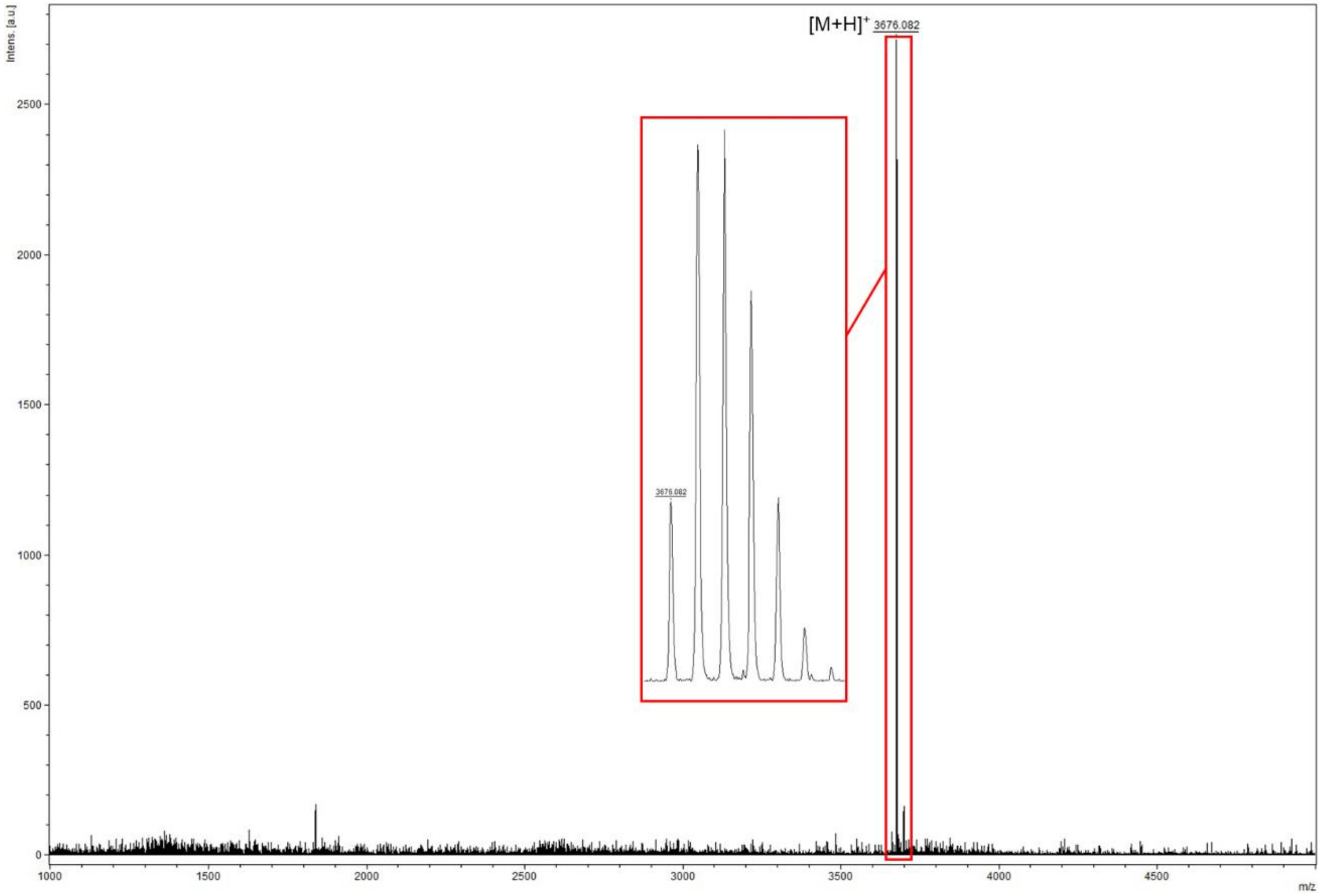


calcd. MS $[M_m + H]^+$: 3675.81 found: 3676.08

Fig. S4. (*Continued*).

**pep4**

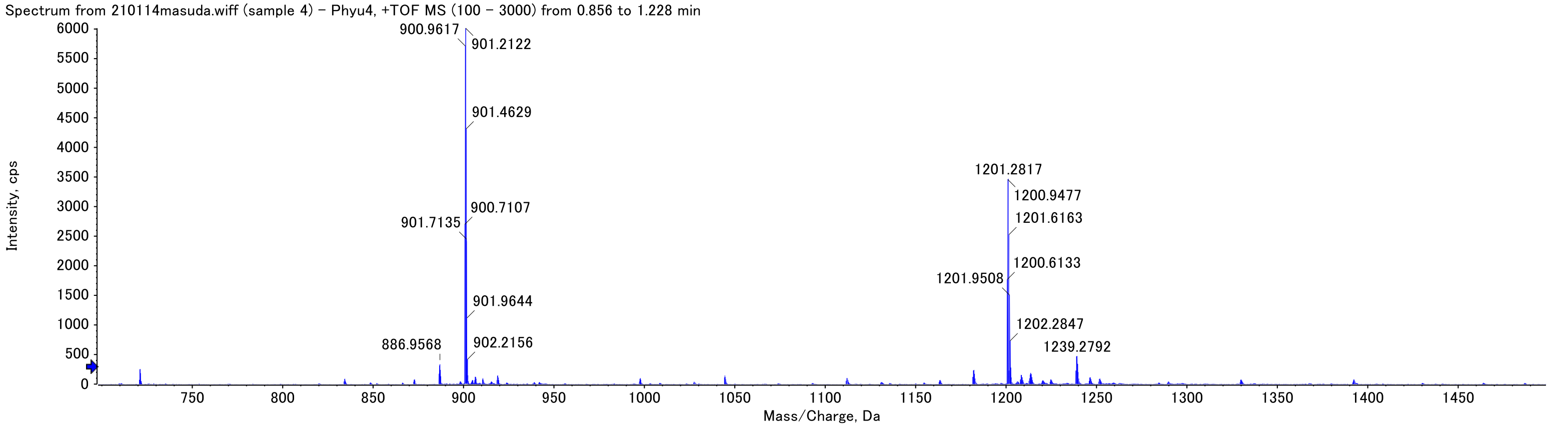


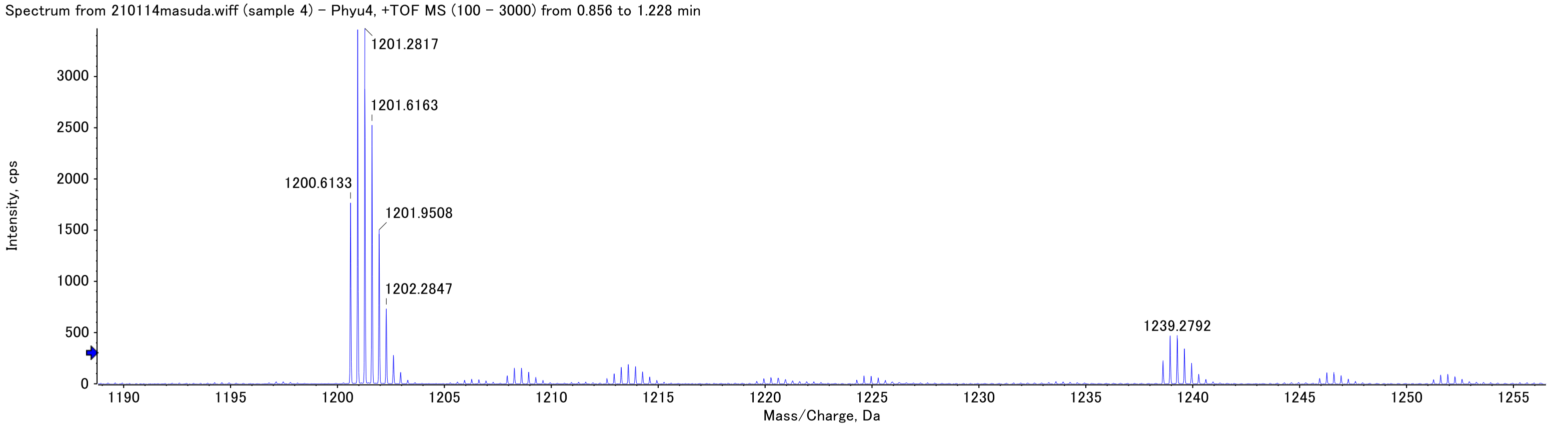


calcd. MS $[M_m + 3H]^{3+}$: 1200.60 found: 1200.61

Fig. S4. (*Continued*).

**pep5**

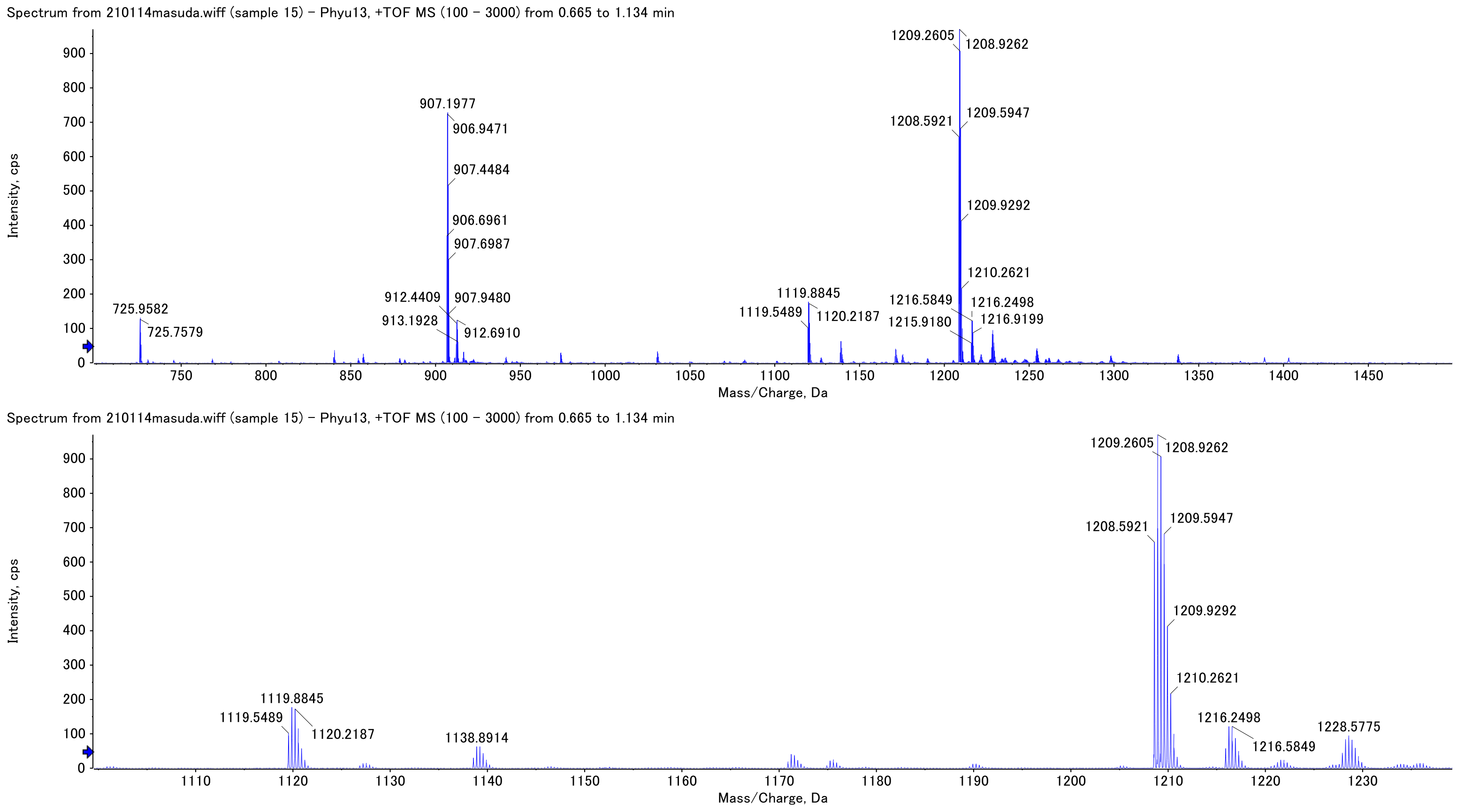


calcd. MS $[M_m + 3H]^{3+}$: 1208.59 found: 1208.59

Fig. S4. (*Continued*).

**pep6**

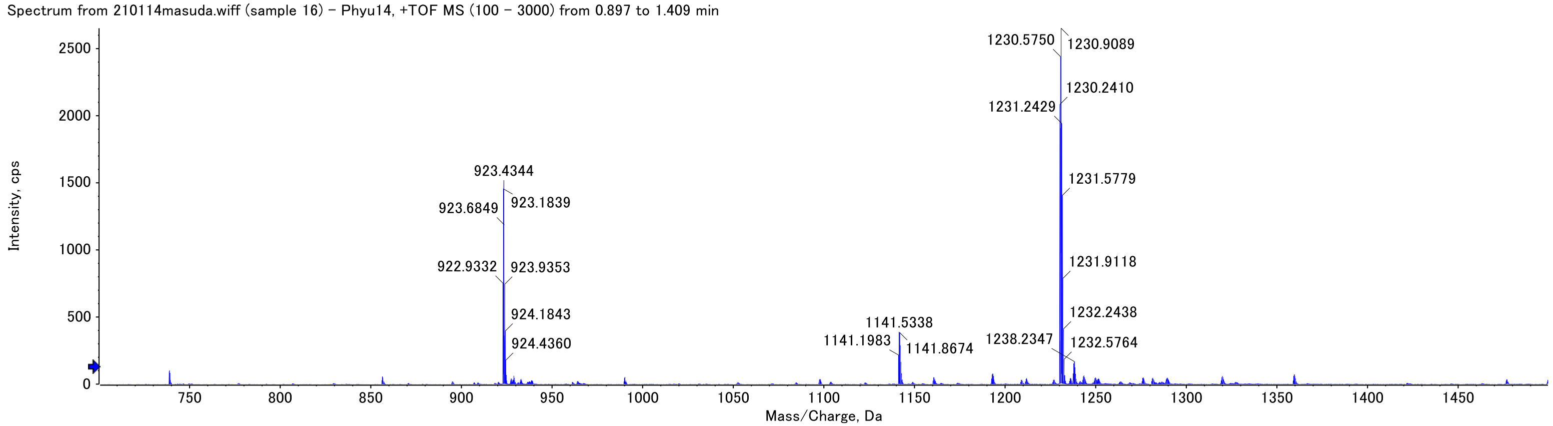


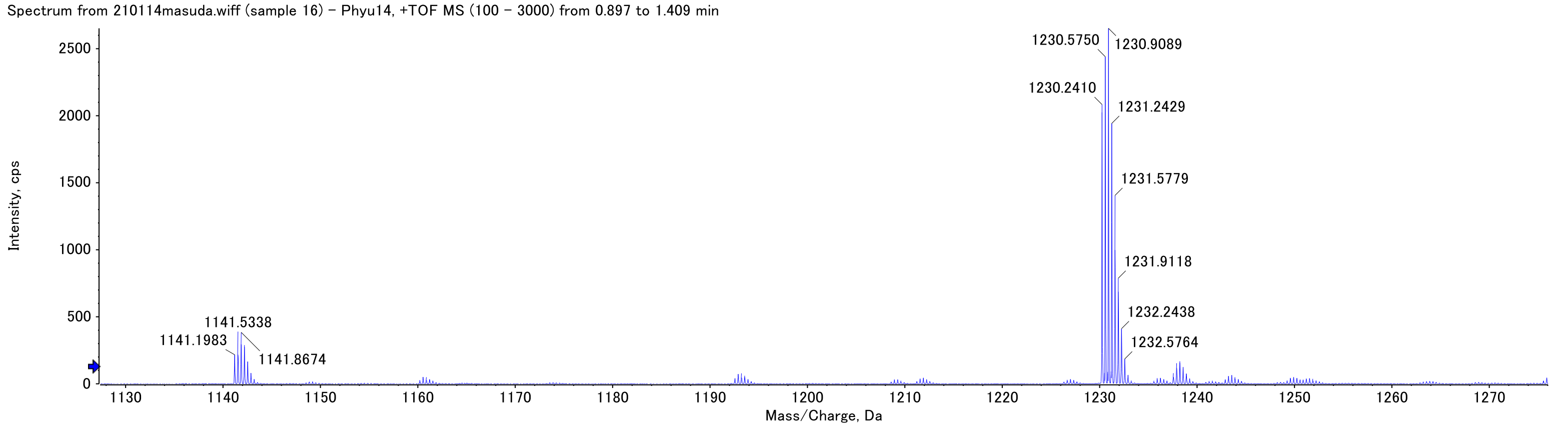


calcd. MS $[M_m + 3H]^{3+}$: 1230.24 found: 1230.24

Fig. S4. (*Continued*).

**pep7**

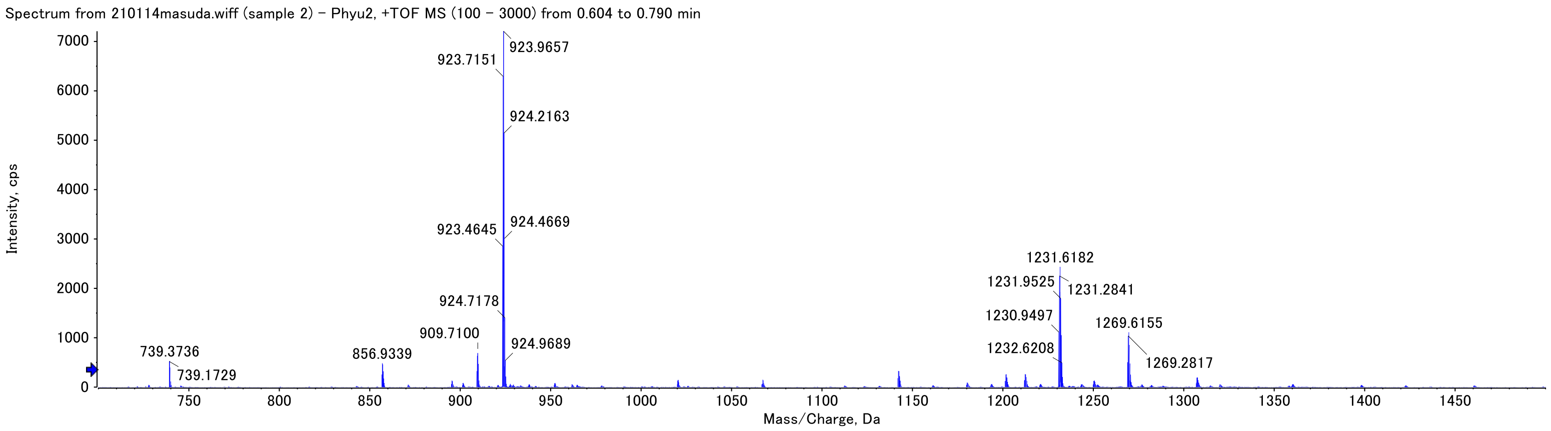


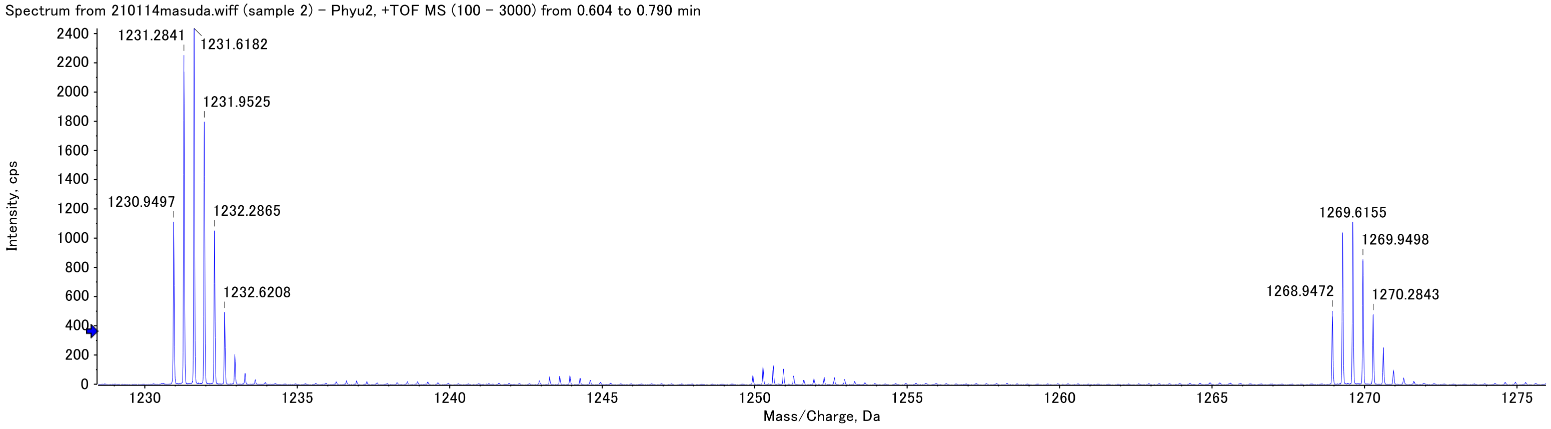


calcd. MS $[M_m + 3H]^{3+}$: 1230.95 found: 1230.95

Fig. S4. (*Continued*).

### pep8

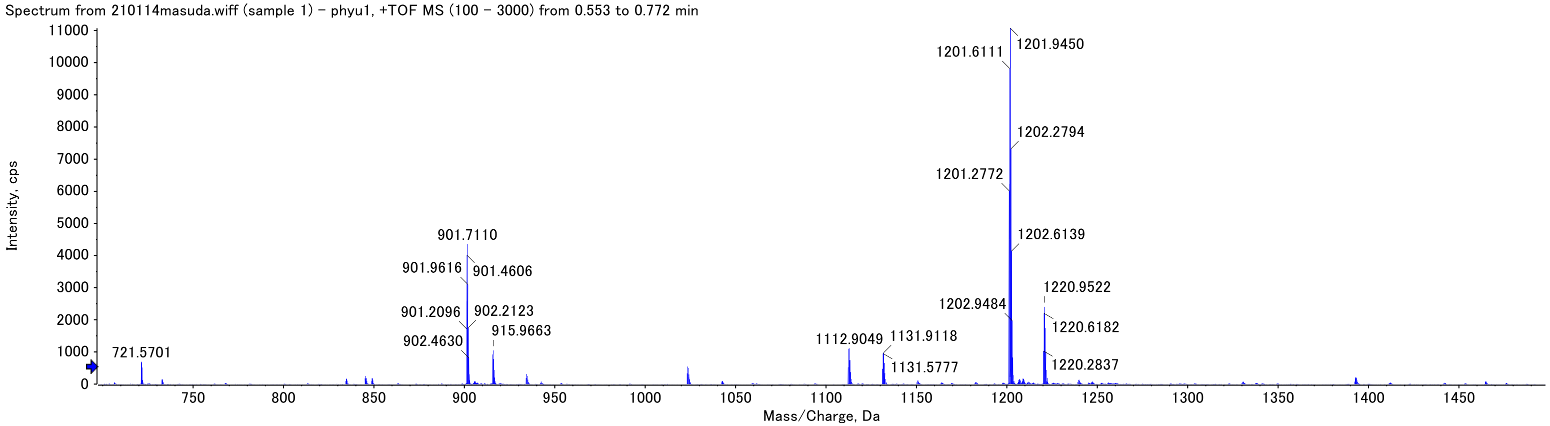


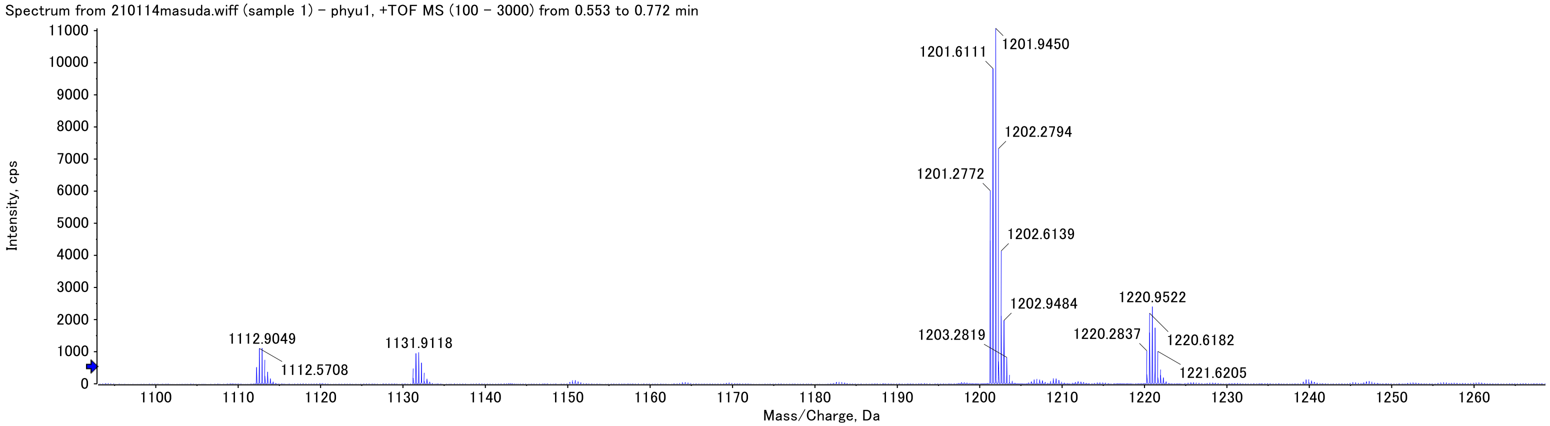


calcd. MS $[M_m + 3H]^{3+}$: 1201.28 found: 1201.28

Fig. S4. (*Continued*).

**pep9**

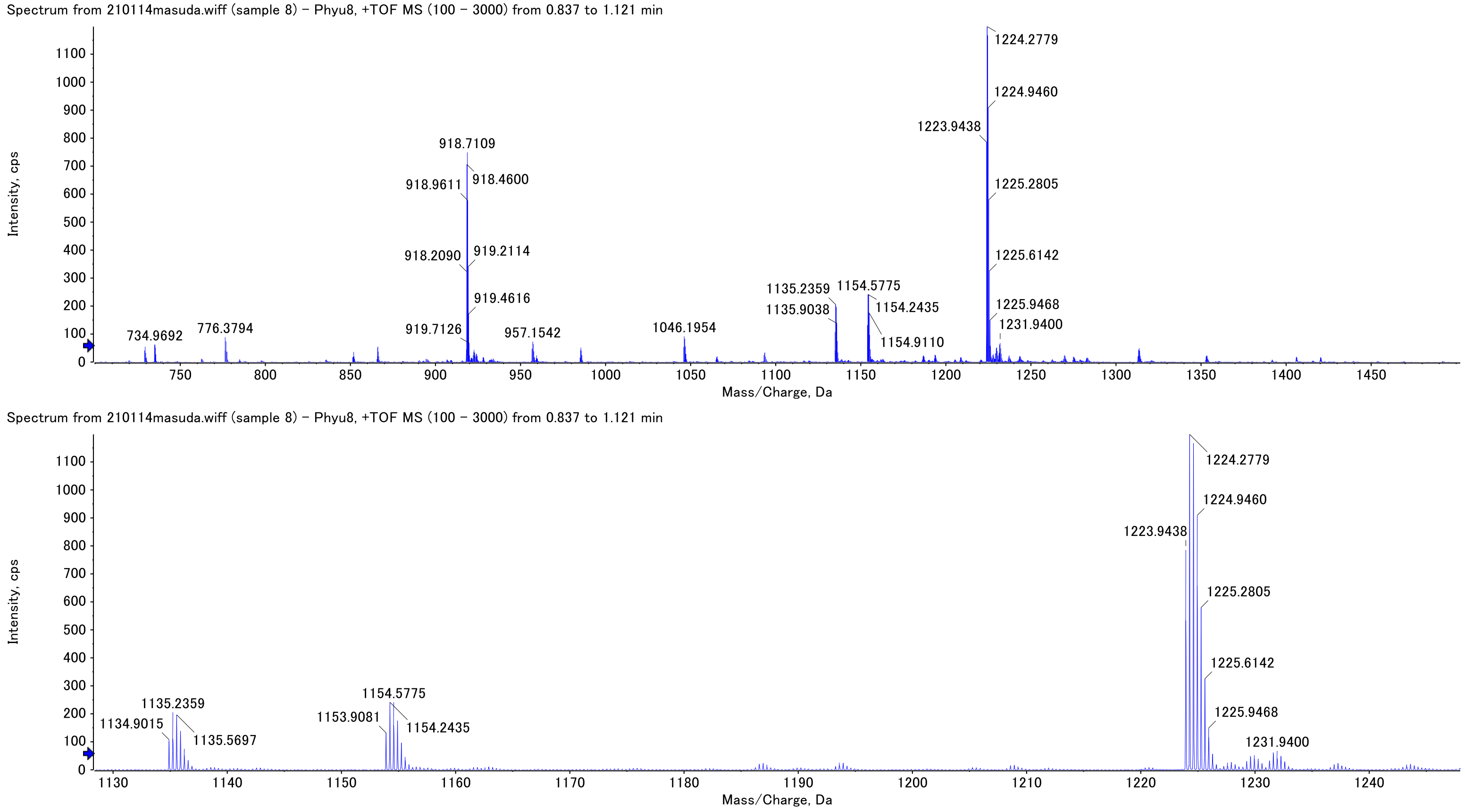


calcd. MS $[M_m + 3H]^{3+}$: 1223.94 found: 1223.94

Fig. S4. (*Continued*).

### pep10

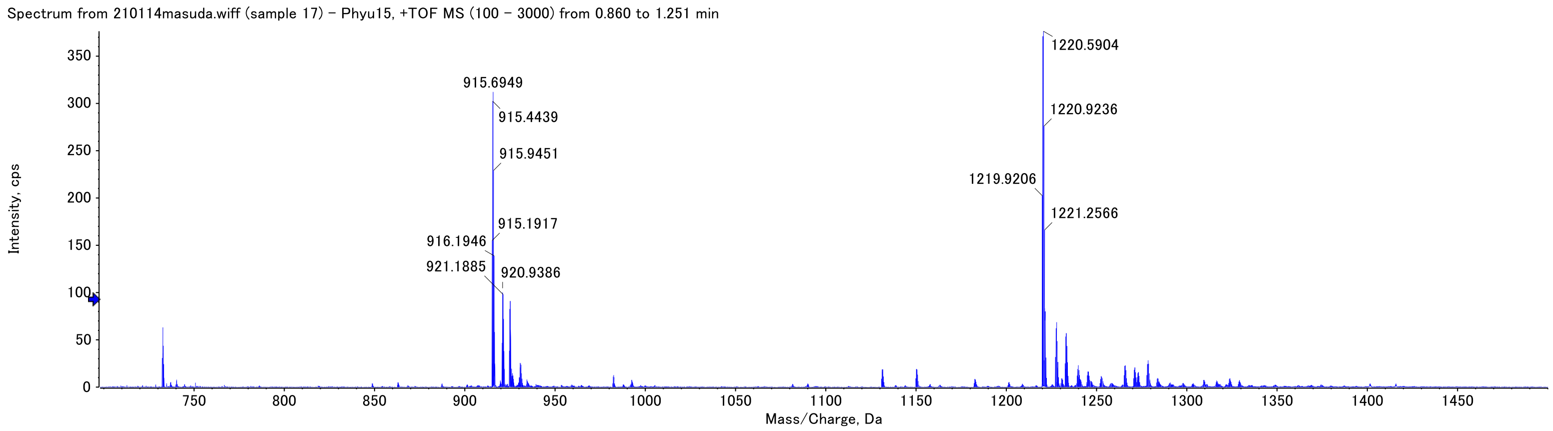


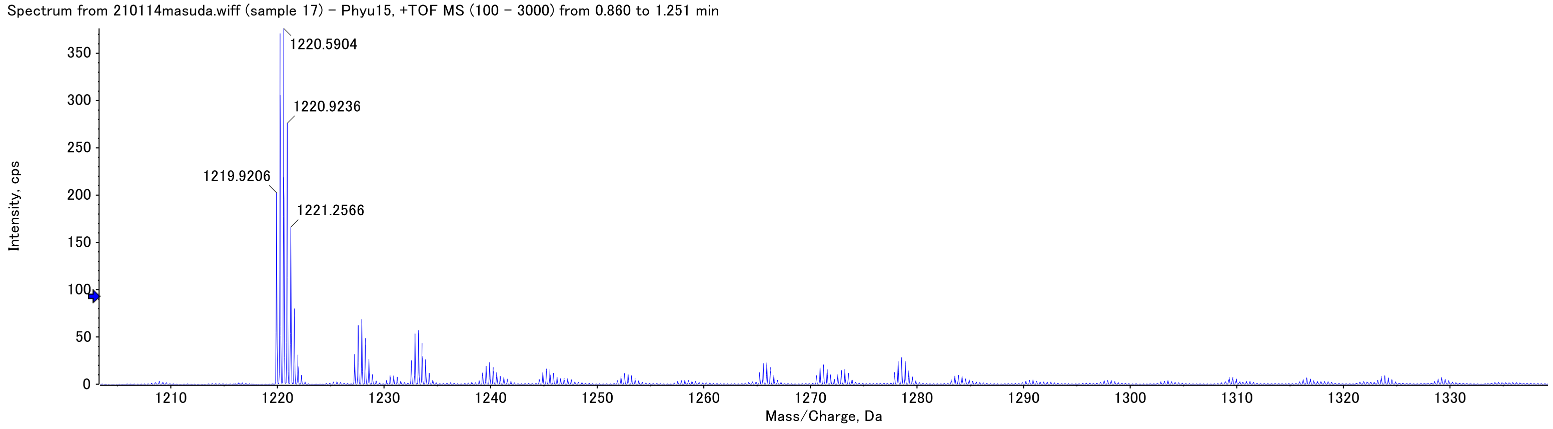


calcd. MS $[M_m + 3H]^{3+}$: 1219.92 found: 1219.92

Fig. S4. (*Continued*).

### pep11

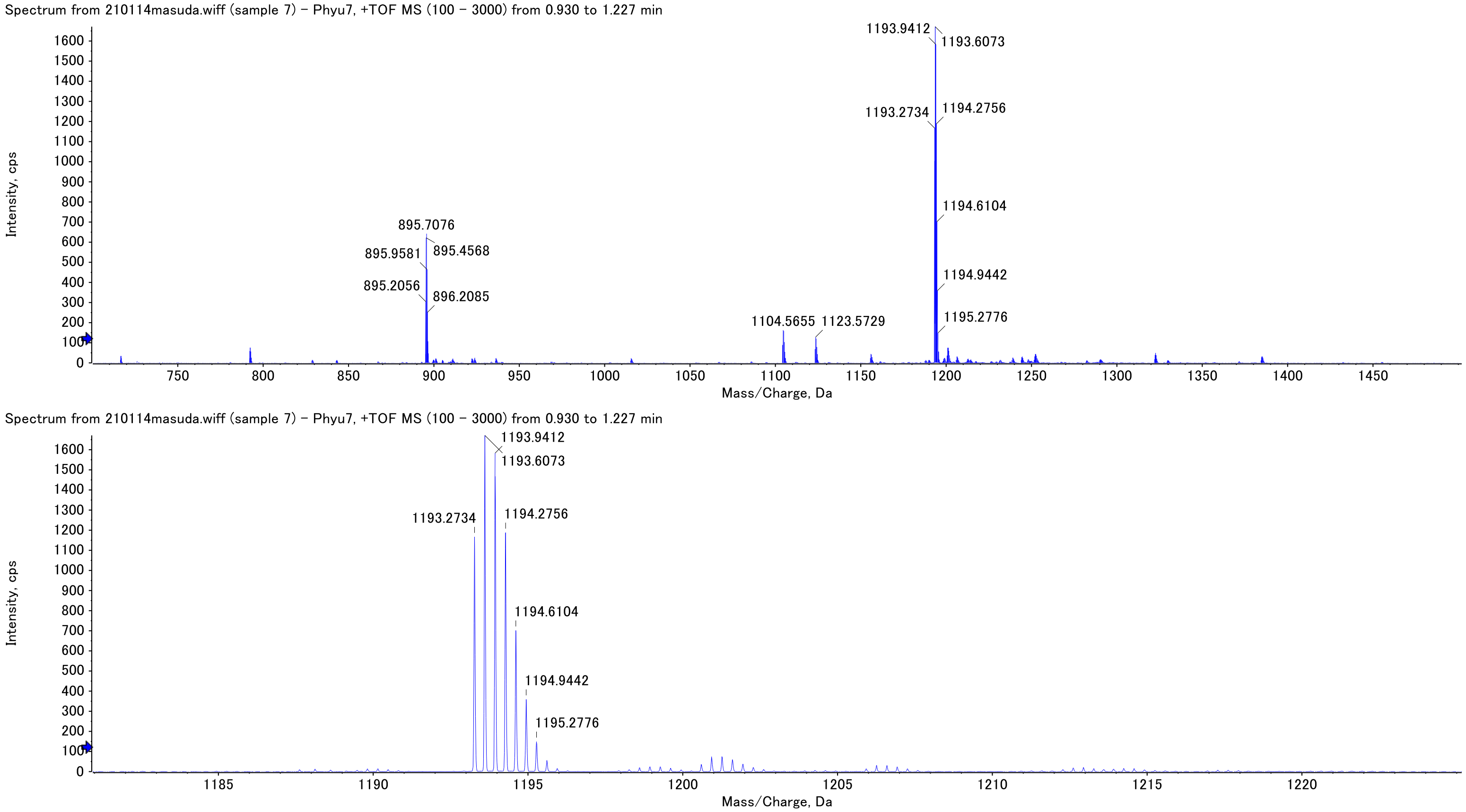


calcd. MS $[M_m + 3H]^{3+}$: 1193.27 found: 1193.27

Fig. S4. (*Continued*).

**pepfreq**

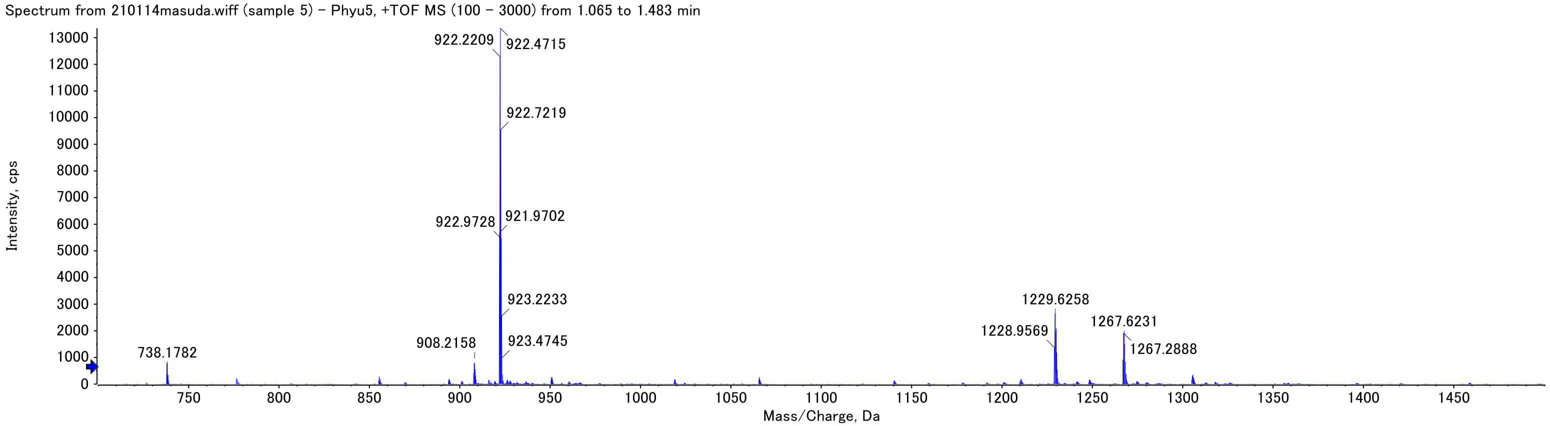


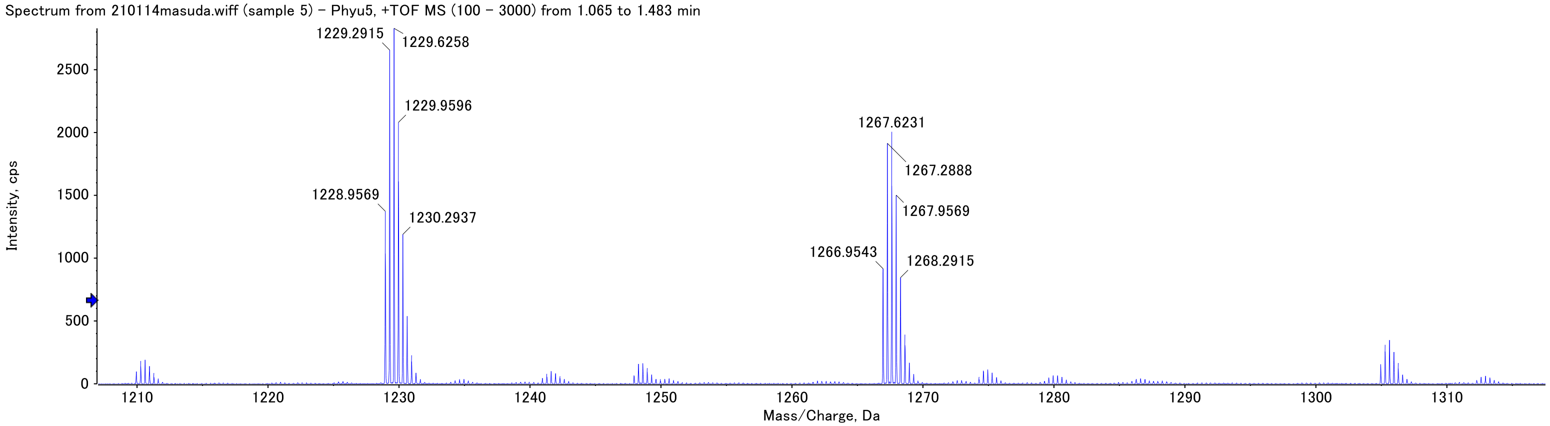


calcd. MS $[M_m + 3H]^{3+}$: 1228.95 found: 1228.96

Fig. S4. (*Continued*).

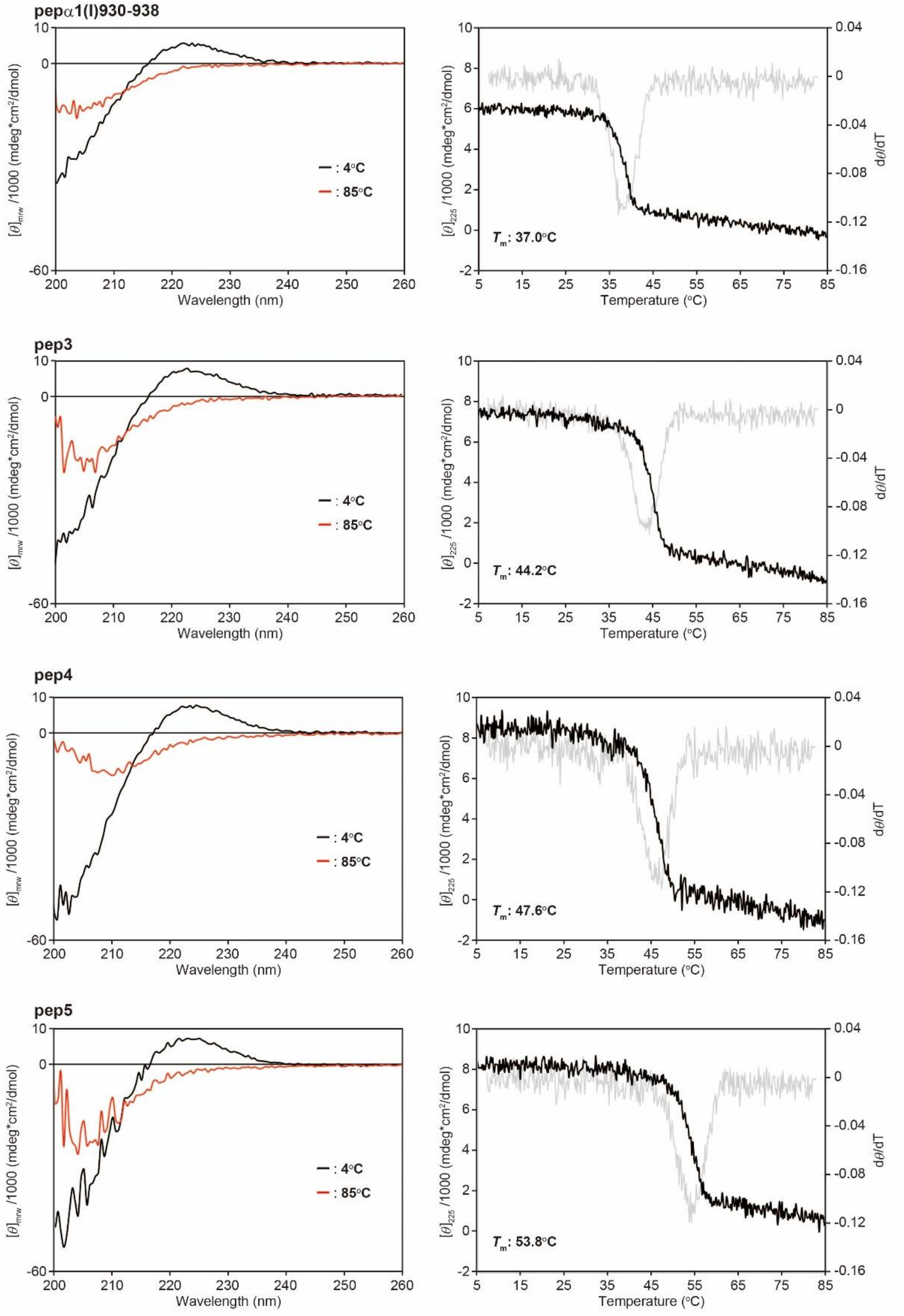


Fig. S5. CD profiles of the synthetic peptides. (Left panels) CD spectrum recorded at 4 and 85°C. (Right panels) A thermal melting curve of the triple helix (black lines) and slope calculated by differentiation (gray lines).

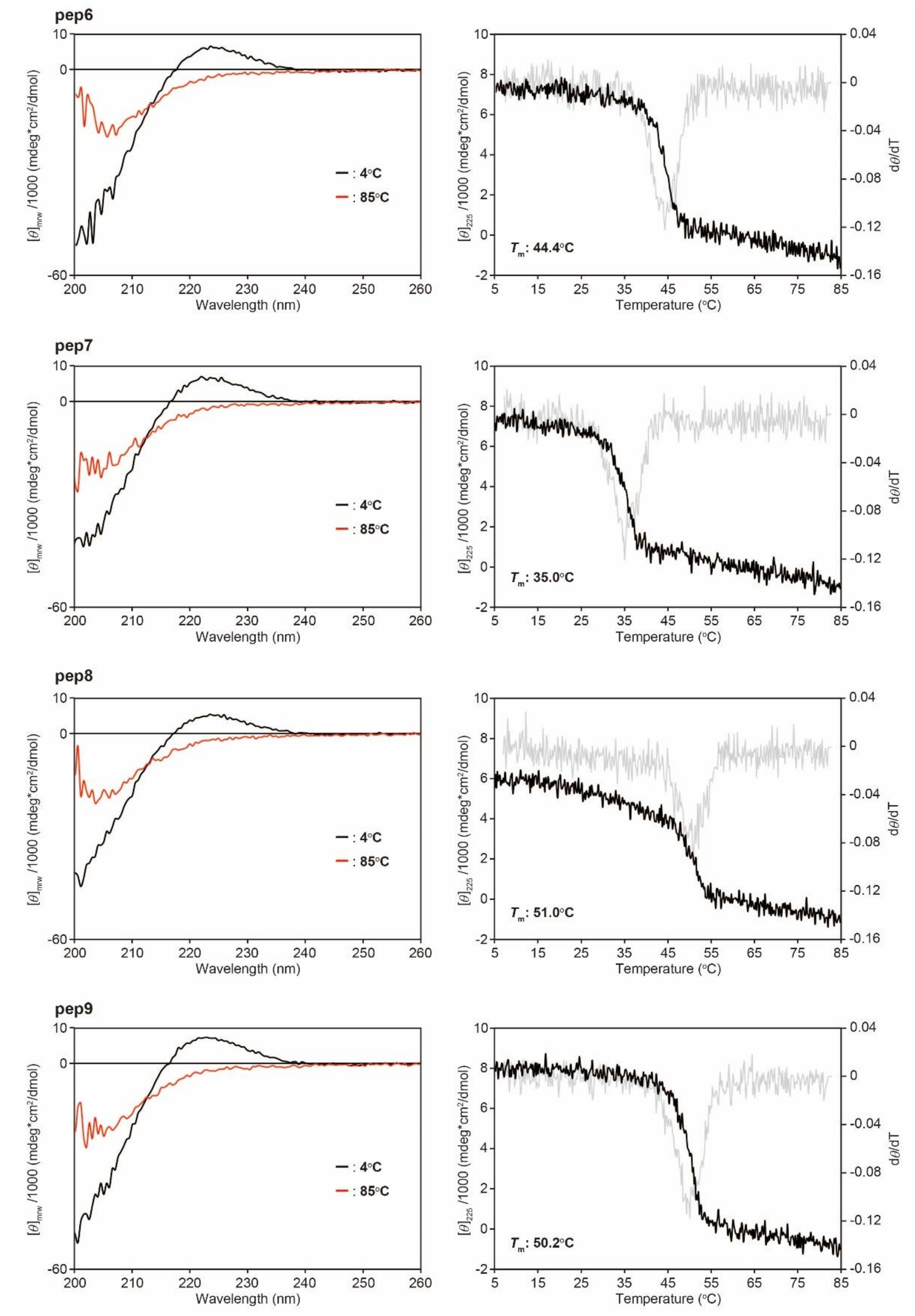


Fig. S5. *(Continued)*.

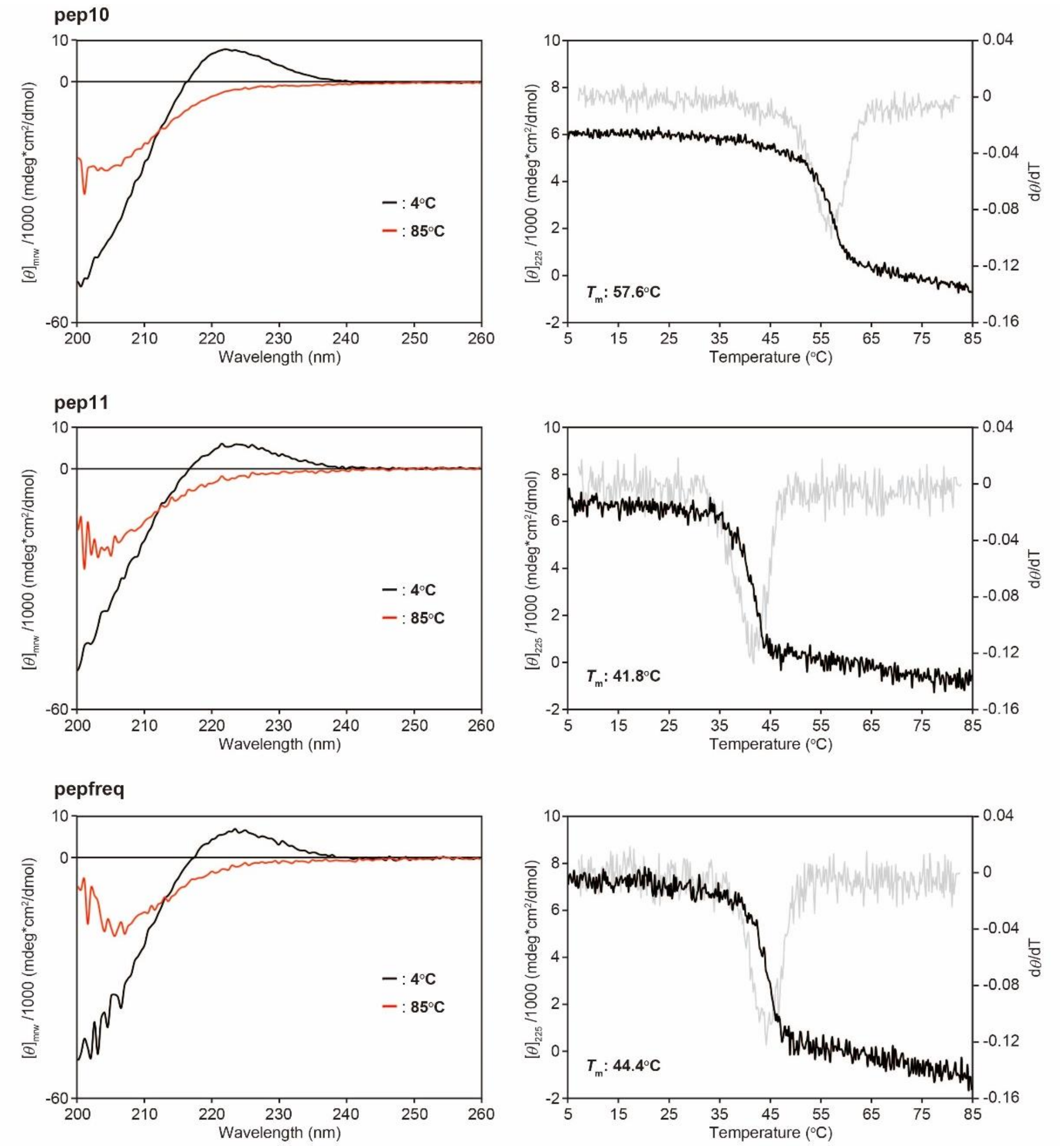


Fig. S5. *(Continued)*.

Table S2. Sequences of primers for preparing the bait constructs used in Fig. 4B

| ssDNA | Sequence |
|---|---|
| PEDF R148A-forward | 5′-AAACTTGCTGTCAAATCCAGCTTTGTT-3′ |
| PEDF R148A-reverse | 5′-TTTGACAGCAAGTTTCCTCTCAAACAC-3′ |
| PEDF D255N-forward | 5′-GGCTTGAATTCTGATCTCAACTGCAAG-3′ |
| PEDF D255N-reverse | 5′-ATCAGAATTCAAGCCGTATCGTAAGAT-3′ |

```
                                      EcoRI
ATG·························GAA TTC CCC GGT CCA CCA GGA CCT CCA GGC CCA
Met·························Glu Phe Pro Gly Pro Pro Gly Pro Pro Gly Pro
        GAL4-AD domain                          (PGP)10

CCC GGT CCG CCT GGT CCT CCC GGT CCT CCC GGA CCA CCA GGC CCA CCT GGG CCG
Pro Gly Pro Pro Gly Pro Pro Gly Pro Pro Gly Pro Pro Gly Pro Pro Gly Pro
                                (PGP)10

       ApaI                                              XmaI
CCA GGG CCC AAA GGT CAT AGA GGT TTT TCT GGT TTG CCC GGG CCA CCG GGT CCA
Pro Gly Pro Lys Gly His Arg Gly Phe Ser Gly Leu Pro Gly Pro Pro Gly Pro
  (PGP)10          PEDF-binding sequence                 (PGP)10

CCC GGA CCT CCA GGT CCG CCA GGC CCT CCT GGC CCT CCG GGT CCA CCA GGT CCA
Pro Gly Pro Pro Gly Pro Pro Gly Pro Pro Gly Pro Pro Gly Pro Pro Gly Pro
                                (PGP)10
                                                        BamHI
CCT GGT CCT CCT GGG CCA CCA GGT TCA GGA TAC ATT TAA GGA TCC
Pro Gly Pro Pro Gly Pro Pro Gly Ser Gly Tyr Ile stop
        (PGP)10
```

Fig. S6 A sequence map of α1(I)930–938/pGADT7

Table S3. Sequences of ssDNA for preparing the prey constructs used in Fig. 3

| ssDNA | Sequence |
|---|---|
| PGP10-sense | 5′-CTGATAGTAATGATAGTAATGATAGTAAC-3′ |
| PGP10-antisense | 5′-CCGGGTTACTATCATTACTATCATTACTATCAGGGCC-3′ |
| PGP13-sense | 5′-CCCAGGTCCTCCAGGTCCTCCAGGTCCTTAGTAAC-3′ |
| PGP13-antisense | 5′-CCGGGTTACTAAGGACCTGGAGGACCTGGAGGACCTGGGGGCC-3′ |
| PGP23-sense | 5′-CCCAGGTCCACCTGGACCACCTGGTCCAC-3′ |
| PGP23-antisense | 5′-CCGGGTGGACCAGGTGGTCCAGGTGGACCTGGGGGCC-3′ |
| 1-1_R4A-sense | 5′-CAAGGGTTGTGCTGGATTGCATGGTCTTC-3′ |
| 1-1_R4A-antisense | 5′-CCGGGAAGACCATGCAATCCAGCACAACCCTTGGGCC-3′ |
| 1-2_R4A-sense | 5′-CAGGGGTGTGGCTGGATTTGAGGGTTGCC-3′ |
| 1-2_R4A-antisense | 5′-CCGGGGCAACCCTCAAATCCAGCCACACCCCTGGGCC-3′ |
| 1-3_R4A-sense | 5′-CCGGGGTCATGCTGGATTTTTGGGTCTTC-3′ |
| 1-3_R4A-antisense | 5′-CCGGGAAGACCCAAAAATCCAGCATGACCCCGGGGCC-3′ |

Table S4. Sequences of primers for NGS samples

| Primer | Sequence |
|---|---|
| NGS-forward | 5′-TCGTCGGCAGCGTCAGATGTGTATAAGAGACAGTACCCATACGACGTACCAGATT-3′ |
| NGS-reverse | 5′-GTCTCGTGGGCTCGGAGATGTGTATAAGAGACAGAGATGGTGCACGATGCACAG-3′ |

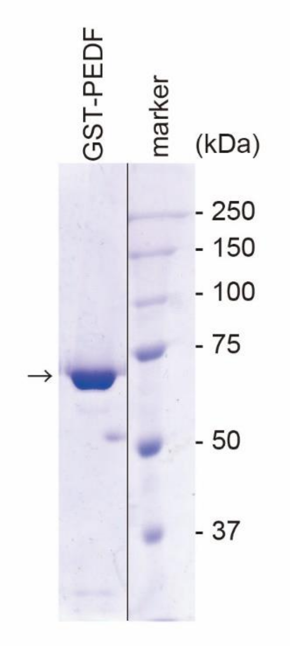


Fig. S7. SDS-PAGE analysis of purified GST-PEDF. Proteins were electrophoresed with 10% acrylamide gel and the bands were visualized by Coomassie Brilliant Blue staining. The molecular sizes are shown in kilodaltons. The molecular weight of GST-PEDF is 72 kDa.
